# Cryptic disease-prone states in human mesocortical assembloids revealed by multimodal profiling

**DOI:** 10.64898/2026.08.01.742201

**Authors:** Seongmin Kim, Rian Kang, Taehoon Lee, Yongjun Kim, Chang-Dae Kim, Kyeong-Mo Koo, Jiyu Na, Sumin Lee, Dohoon Kim, Hayoung Oh, Amos Chungwon Lee, Tae-Hyung Kim, Byullee Park, Luke P. Lee, Inki Kim, Jong-Chan Park

## Abstract

Neurodegenerative diseases are characterized by synaptic failure, aberrant protein accumulation, and neuroglial dysfunction that emerge long before clinical onset. Although brain assembloids, which recapitulate interregional circuit connectivity beyond the scope of single organoids, significantly advance the modeling of circuit pathophysiology, they are predominantly evaluated through single-modality approaches that cannot resolve the functional and molecular heterogeneity underlying differential disease susceptibility. Here we show that morphologically identical human iPSC-derived dorso forebrain-midbrain mesocortical assembloids (MCAs) spontaneously bifurcate into disease-prone and non-prone states under identical culture conditions, revealing that MCA heterogeneity reflects intrinsic neurodegeneration susceptibility rather than stochastic culture variability. Using integrated electrophysiological, molecular, and spatial profiling, we find that disease-prone MCAs exhibit a temporally ordered molecular cascade in which neurofilament light chain elevation precedes tau dysregulation, mirroring the sequential biomarker trajectories observed in pre-symptomatic human neurodegeneration. Disease-prone MCAs further display selective cortical hyperexcitability and aberrant brainwave-like oscillatory dynamics that remain undetectable by any single modality. Spatially, a discrete junction-like neuronal population at the midbrain-forebrain interface shows transcriptional priming for synaptic overactivation alongside impaired astrocytic glutamate clearance, defining a spatially confined neuron-glial uncoupling as a candidate early origin of the disease-prone state. These findings reframe MCA heterogeneity as a biological window into pre-symptomatic neurodegeneration, with broad implications for disease modeling, risk stratification, and therapeutic discovery.

## Introduction

Understanding the mechanisms underlying neurodegeneration is a priority in neuroscience research given its profound impact on cognitive function, neuronal integrity, and brain homeostasis. Neurodegeneration is characterized by progressive neuronal dysfunction, disrupted neuron-glial interactions^1^, metabolic dysregulation^2^, and pathological protein accumulation^3^. However, traditional approaches for studying neurodegeneration predominantly rely on single-modality assessments, such as endpoint histological staining, isolated electrophysiological recordings, and destructive biochemical assays, that cannot capture the spatial heterogeneity^4^ and temporal dynamics underlying disease progression, thus limiting our ability to predict and characterize disease-prone states before irreversible damage.

Recent advances in brain organoid technology have significantly improved the ability to model human brain development and neurodegeneration in vitro^5–7^. Brain assembloids that physically fuse multiple organoid regions, such as forebrain (FO) and midbrain-like (MO) organoids, provide a valuable opportunity to study interregional connectivity^8^ and circuit-level pathophysiology^9^ systematically. The FO–MO assembloid specifically recapitulates the corticodopaminergic pathway, a critical circuit implicated in Parkinson’s disease^10,11^, schizophrenia^12,13^, and autism spectrum disorders^14–16^. However, substantial inter-organoid variability in functional and molecular signatures has emerged as a fundamental challenge even under identical genetic backgrounds and culture conditions^17^. This heterogeneity has traditionally been considered a technical limitation requiring standardization. We propose, however, that intrinsic variability in assembloid phenotype may reflect biologically meaningful differences in disease susceptibility, mirroring the inter-individual variation in neurodegeneration vulnerability observed in human populations.

To test this hypothesis, we simultaneously interrogated neural circuit dynamics, molecular pathology, and spatial transcriptional architecture across human iPSC-derived dorso forebrain-midbrain mesocortical assembloids (MCAs) — integrating HD-MEA electrophysiology^18,19^, immuno-SERS-based detection of T-tau, p-tau, NfL, and CD81^20–22^, and spatially resolved transcriptomics via laser-activated cell sorting (SLACS). This integrated strategy uncovered a previously unrecognized biological phenomenon — morphologically identical MCAs spontaneously bifurcate into disease-prone and non-prone states under identical culture conditions, a divergence whose full biological depth, spanning transcriptional reprogramming, protein secretion dynamics, bioenergetic state, and circuit-level electrophysiology, is resolvable only through cross-modal integration of these complementary analytical dimensions. Inter-MCA variability in circuit excitability, protein secretion, and transcriptional state reflects intrinsic differences in neurodegeneration susceptibility rather than stochastic culture variability, mirroring the inter-individual heterogeneity observed in human neurodegenerative risk. Disease-prone MCAs are defined by spontaneous cortical hyperexcitability, a temporally ordered cascade of neurofilament and tau accumulation, and convergent multimodal signatures that together enable prospective risk classification — with spatial transcriptomic analysis further revealing region-specific neuron-glial uncoupling at the forebrain-midbrain junction as a candidate mechanistic origin of the disease-prone state.

## Results

### Morphological uniformity of dorso forebrain-midbrain mesocortical assembloids (MCAs) does not capture the full spectrum of functional and molecular phenotypes

To model the human mesocortical neuronal circuit, we generated dorso forebrain-midbrain mesocortical assembloids (MCAs) by fusing human iPSC-derived forebrain organoids (FOs) and midbrain-like organoids (MOs) (**Fig. 1a**). FOs underwent sequential ectoderm induction, neural differentiation (bFGF/EGF), and maturation (BDNF/NT3) over 100 days. MOs were independently patterned through midbrain specification (IM−, IM+, TG+, OG+) over 35 days. Fusion of day-100 FOs and MOs was performed in tilted 24-well plates to facilitate initial cell–cell adhesion, followed by orbital shaker culture for two weeks. Fluorescence imaging confirmed robust structural integration between green fluorescent protein (GFP)-labeled FO and red fluorescent protein (RFP)-labeled MO compartments, with neurons in the MO region projecting axons toward the FO region (**Fig. 1b**), recapitulating the mesocortical circuitry implicated in Parkinson’s disease, schizophrenia, and autism spectrum disorders. Regional characterization of each organoid was further validated by immunohistochemistry, showing selective FOXG1 expression in the FO and tyrosine hydroxylase (TH) enrichment in the MO, confirming successful regional specification of MCAs (**Extended Data fig. 1**).

**Fig. 1.**
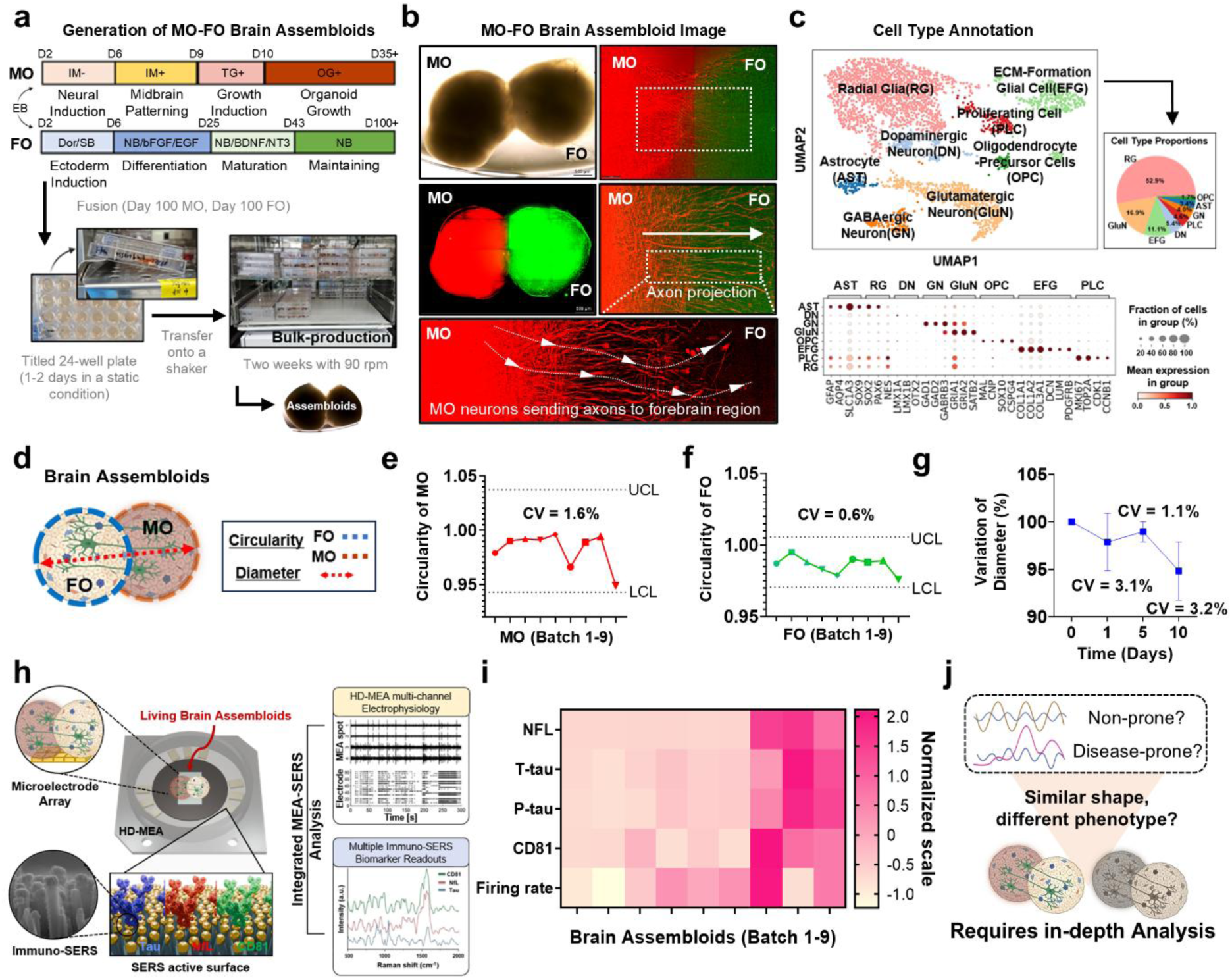
Morphological uniformity of dorso forebrain-midbrain mesocortical assembloids (MCAs) does not capture the full spectrum of functional and molecular phenotypes. **(a)** Differentiation protocols for forebrain organoids (FO) and midbrain-like organoids (MO) from human iPSCs, showing sequential induction stages and media compositions. MCA bulk-production via tilted 24-well plate fusion followed by orbital shaker (90 rpm) culture. **(b)** Representative bright-field and dual-fluorescence images of MCAs showing structural integration of GFP-labeled FO and RFP-labeled MO, with axonal projections from MO to FO region. **(c)** UMAP visualization of scRNA-seq data with cell-type annotation of major neural and glial populations, dot plot of canonical marker gene expression, and proportional cell-type composition. **(d)** Schematic morphological parameters to quantify circularity and diameter of MCAs. **(e, f)** Circularity of FO (CV = 0.6%) and MO (CV = 1.6%) across Batch 1–9. UCL, upper control limit; LCL, lower control limit. **(g)** Diameter variation (%) over 10 days across assembloid groups, with inter-batch CV values indicated. **(h)** Schematic of the integrated HD-MEA and Immuno-SERS platform for simultaneous electrophysiological recording and secreted protein detection (T-tau, NfL, CD81). **(i)** Normalized heatmap of NfL, T-tau, p-tau, CD81, and MEA-derived firing rate across MCAs from Batch 1–9. **(j)** Schematic illustrating that morphologically equivalent MCAs can harbor functionally distinct disease-prone (DP) or non-prone (NP) phenotypes, necessitating in-depth multimodal analysis.

Single-cell RNA sequencing confirmed the expected cellular composition across MCAs (**Fig. 1c**). UMAP-based clustering identified eight major cell populations with distinct transcriptional identities. Radial glia (RG; 52.9%), the most abundant population, were marked by *PAX6* and *SOX2*. Glutamatergic neurons (GluN; 16.9%) expressed *GRIA1*, *GRIA2*, and *SATB2*, while GABAergic neurons (GN; 4.0%) were identified by *GAD1*, *GAD2*, and *GABRB3*. Dopaminergic neurons (DN; 5.4%) were characterized by midbrain-specific transcription factors *LMX1A*, *LMX1B*, and *OTX2*. Astrocytes (AST; 3.4%) expressed *GFAP* and *AQP4*, and Oligodendrocyte-Precursor Cells (OPC; 1.7%) were marked by *SOX10*, *CSPG4*, *MAL*, and *CNP*. ECM-Formation Glial Cells (EFG) were distinguished by *COL1A1*, *DCN*, and *LUM*, and proliferating cells (PLC; 4.6%) expressed cell cycle markers *MKI67*, *TOP2A*, and *CDK1*. Dot plot analysis confirmed selective enrichment of these canonical marker genes within each annotated cluster, validating the cellular diversity and regional identity of the MCA model. Visualization of individual marker expression on UMAP plots further validated the distinct transcriptional identities of the annotated neuronal and glial populations (**Extended Data fig. 2**).

In addition, morphological reproducibility was systematically assessed across nine independent MCA batches (**Fig. 1d-g**). Circularity of MO and FO components remained stable throughout production (CV = 1.6% and 0.6%, respectively; **Fig. 1e, f**), and assembloid diameter showed consistent growth trajectories over 10 days with low inter-batch variance (CV = 1.1–3.2%; **Fig. 1g**). For multimodal profiling, MCAs were interfaced with a high-density microelectrode array (HD-MEA) platform co-localized with immuno-SERS substrates enabling simultaneous electrophysiological recording and multiplexed detection of secreted neurodegeneration markers—T-tau, NfL, and CD81 (**Fig. 1h**). The immuno-SERS substrates were fabricated using antibody-functionalized ZnO nanowire arrays optimized for sensitive multiplexed protein detection (**Extended Data fig. 3**). Despite similar morphological uniformity, normalized multimodal profiles across Batch 1–9 revealed substantial inter-assembloid variation in NfL, T-tau, p-tau, CD81, and MEA-derived firing rate (**Fig. 1i**), demonstrating that morphological metrics are insufficient to predict functional or molecular states. These findings establish that morphological equivalence does not reflect phenotypic equivalence in MCAs — functional and molecular divergence between disease-prone and non-prone populations remains invisible to morphological assessment alone, underscoring the necessity of multimodal characterization (**Fig. 1j**).

### SERS-based molecular risk profiling coupled with MEA characterization identifies disease-prone and non-prone MCAs

To systematically classify the phenotypic heterogeneity observed across MCA batches, we performed integrated molecular and electrophysiological profiling combining immuno-SERS-based marker quantification with HD-MEA-based network analysis (**Fig. 2**). Classification thresholds were defined based on the upper quartile of normalized marker intensities across all assembloids, ensuring that designations reflect genuine distributional extremes rather than arbitrary cutoffs. MCAs were designated “disease-prone” when normalized SERS intensities exceeded thresholds for at least two markers (T-tau > 1.0, p-tau > 0.4, or NfL > 0.5), a multi-marker criterion adopted to minimize false classification driven by single-marker variability. MCAs maintaining all parameters below threshold were designated “non-prone” (**Fig. 2a**). Electrochemical ATP sensing revealed significantly elevated bioenergetic activity in disease-prone MCAs, with disease-prone assembloids displaying higher ATP release compared with non-prone controls (**Extended Data fig. 4**), indicating that mitochondrial metabolic dysregulation accompanies the molecular and electrophysiological signatures of the disease-prone state. Longitudinal immuno-SERS monitoring revealed a temporally structured progression of molecular pathology in disease-prone MCAs (**Fig. 2b**). At 6 hours, T-tau, p-tau, and NfL levels were indistinguishable between groups. By 24 hours, NfL emerged as the earliest diverging marker, showing significant elevation in disease-prone MCAs (*P* < 0.001), while T-tau and p-tau remained comparable between groups. By 48 hours, all three markers were significantly elevated in disease-prone MCAs (T-tau: *P* < 0.01; p-tau: *P* < 0.001; NfL: *P* < 0.0001), defining a progressive molecular pathology window in which NfL serves as the earliest detectable indicator of neurodegeneration.

**Fig. 2.**
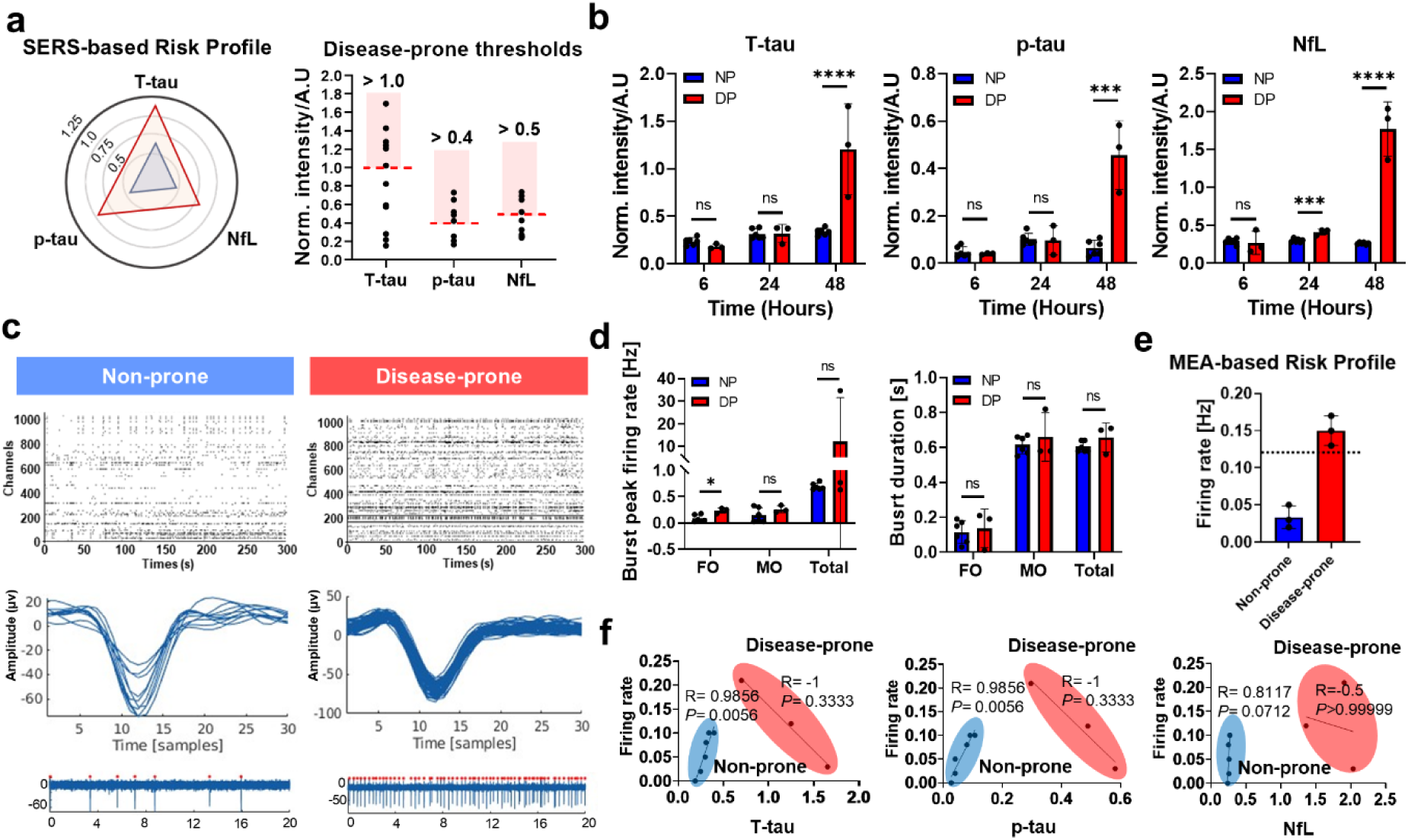
SERS-based molecular risk profiling coupled with MEA characterization identifies disease-prone and non-prone MCAs. **(a)** SERS-based risk profile radar plots of normalized T-tau, p-tau, and NfL intensities, along with scatter plots defining the disease-prone thresholds for each biomarker. **(b)** Time-course quantification of T-tau, p-tau, and NfL levels by immuno-SERS at 6, 24, and 48 h. Statistical significance: ns, not significant; ** *P* < 0.01; *** *P* < 0.001; **** *P* < 0.0001. **(c)** Representative HD-MEA raster plots, averaged spike waveforms, and raw extracellular voltage traces from non-prone and disease-prone assembloids. **(d)** Quantification of burst peak firing rate and burst duration in total, FO, MO, and total regions for non-prone and disease-prone MCAs. Statistical significance: ns, not significant; * *P* < 0.05. **(e)** MEA-based risk profile bar graph comparing spontaneous firing rate between non-prone and disease-prone MCAs. **(f)** Correlation plots between spontaneous firing rate and SERS-derived marker intensities (NfL, T-tau, and p-tau) for non-prone and disease-prone MCAs, with Spearman correlation coefficients (R) and P-values indicated.

HD-MEA recordings revealed clear electrophysiological differences between the two groups (**Fig. 2c**), further supported by representative multichannel recordings, spike waveforms, and spatial electrode maps showing distinct firing dynamics in non-prone and disease-prone MCAs (**Extended Data fig. 5**). Non-prone MCAs displayed regular, coordinated burst patterns with consistent interburst intervals. In contrast, disease-prone MCAs exhibited disorganized bursting with prolonged silent periods and desynchronized firing, consistent with a pattern of network hyperexcitability interspersed with periods of suppressed activity. Region-specific analysis showed that while total and MO-region burst metrics were comparable between groups, disease-prone MCAs exhibited a significant elevation in burst peak firing rate specifically in the FO region (*P* < 0.05; **Fig. 2d**), indicating localized cortical hyperexcitability rather than global network disruption. Corroborating this finding, spontaneous firing rate was significantly elevated in disease-prone MCAs (**Fig. 2e**), consistent with the SERS-based classification and further supporting spontaneous firing rate as a defining functional feature of the disease-prone phenotype.

Correlation analysis between spontaneous firing rate and SERS-derived marker intensities revealed fundamentally distinct coupling patterns between the two populations (**Fig. 2f**). Non-prone MCAs showed positive Spearman correlations between firing rate and T-tau (R = 0.9856, *P* < 0.01) and p-tau (R = 0.9856, *P* < 0.01), while NfL did not reach significance (R = 0.8117, *P* = 0.0712). In contrast, disease-prone MCAs exhibited absent or negative correlations across all three markers—NfL (R = −0.5, *P* > 0.99), T-tau (R = −1, *P* = 0.3333), and p-tau (R = −1, *P* = 0.3333). These divergent correlation structures suggest that the coordinated relationship between network excitability and protein marker levels observed in non-prone MCAs is fundamentally disrupted in the disease-prone state. The exosomal marker CD81 progressively increased in disease-prone MCAs and showed strong correlations with both electrophysiological parameters and SERS-derived neurodegeneration markers, implicating extracellular vesicle dynamics as a late-stage integrative component of disease-prone progression (**Extended Data fig. 6**).

### Disease-prone MCAs display aberrant brainwave-like oscillatory dynamics and are quantified by a PCA-based composite risk score

To characterize network-level oscillatory dynamics beyond burst-level metrics, we performed spectral analysis of HD-MEA-derived local field potentials (LFPs). Raw broadband signals were processed through a 300 Hz high-pass filter, and LFPs were decomposed into delta, theta, alpha, beta, and gamma frequency bands via fast Fourier transform (FFT) (**Fig. 3a**). This frequency-domain framework parallels clinical electroencephalography (EEG) analysis, enabling translational comparison with disease-associated brainwave signatures.

**Fig. 3.**
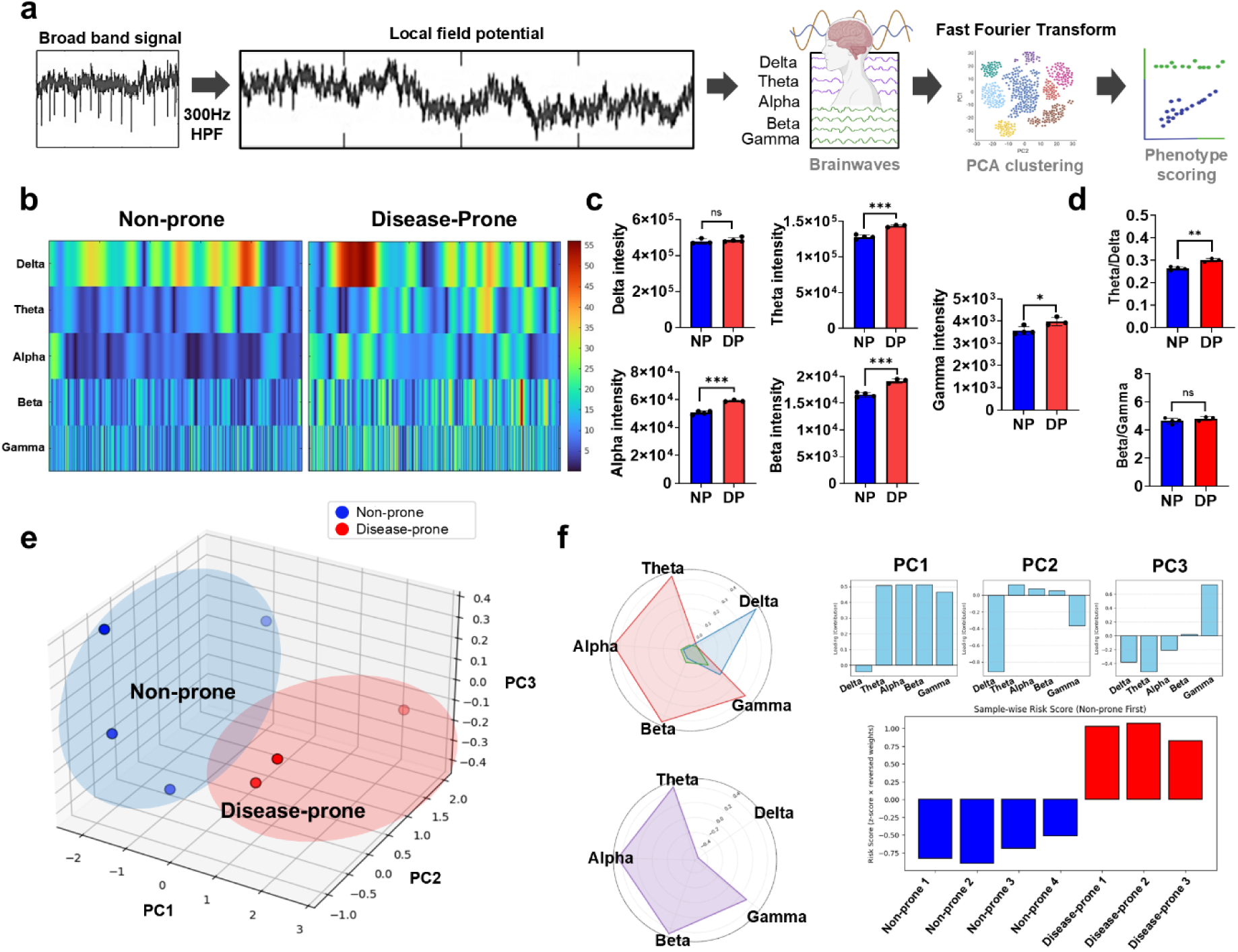
Disease-prone MCAs display aberrant brainwave-like oscillatory dynamics and are quantified by a PCA-based composite risk score. **(a)** Analysis pipeline from raw broadband MEA signal through 300 Hz high-pass filtering for LFP isolation, FFT-based decomposition into delta, theta, alpha, beta, and gamma frequency bands, PCA-based clustering, and composite phenotype risk scoring. **(b)** Functional network connectivity heatmaps across frequency bands over time for non-prone and disease-prone MCAs. **(c)** Quantitative comparison of delta, theta, alpha, beta, and gamma band power between non-prone and disease-prone MCAs. **(d)** Theta/delta and beta/gamma ratios between non-prone and disease-prone MCAs. Statistical significance: ns, not significant; * *P* < 0.05; ** *P* < 0.01; *** *P* < 0.001. **(e)** Three-dimensional PCA scatter plot of multiband oscillatory features for non-prone and disease-prone MCAs. **(f)** Radar plots and bar graphs of PC1, PC2, and PC3 frequency-band loading contributions for non-prone and disease-prone MCAs, and PCA-based composite risk scores per individual MCA sample.

Functional network connectivity heatmaps confirmed organized low-frequency temporal dynamics in non-prone MCAs versus chaotic spectrotemporal patterns with disrupted delta coherence in disease-prone MCAs (**Fig. 3b**). Quantitative band power analysis confirmed significant group differences across multiple frequency bands (**Fig. 3c**). While delta-band power did not differ significantly between groups, disease-prone MCAs showed significantly elevated theta (*P* < 0.001), alpha (*P* < 0.001), beta (*P* < 0.001), and gamma (*P* < 0.05) band power. The theta/delta ratio was significantly elevated in disease-prone MCAs (*P* < 0.01; **Fig. 3d**), reflecting a relative shift toward higher-frequency oscillatory activity. This pattern parallels the theta/delta ratio increases reported in EEG studies of Alzheimer’s disease, suggesting that MCAs may recapitulate aspects of disease-associated network dynamics at the frequency-domain level. In contrast, the beta/gamma ratio did not differ significantly between groups (**Fig. 3d**), indicating that the relative balance between high-frequency bands remains preserved despite their overall elevation.

Principal component analysis (PCA) of multiband oscillatory features separated NP and DP populations in three-dimensional feature space (**Fig. 3e**). Analysis of PC loading contributions revealed that theta and alpha band activities were the primary drivers of the disease-prone oscillatory signature, while delta activity predominated in the non-prone cluster (**Fig. 3f**). A PCA-weighted composite risk score consistently assigned negative scores to non-prone MCAs and positive scores to disease-prone MCAs across all tested assembloids, providing a robust and interpretable quantitative readout for phenotype classification (**Fig. 3f**).

### Integrated multimodal analysis identifies disease-prone risk signatures in assembloids

Having established distinct electrophysiological, oscillatory, and molecular signatures of disease-prone MCAs across independent modalities, we next asked whether integrating these complementary modalities could reveal cross-modal relationships, linking transcriptional programs identified by spatial analysis to protein secretion dynamics, bioenergetic state, and circuit-level electrophysiology, that are inaccessible when each dimension is considered in isolation. To address this, we performed unsupervised multimodal integration of the features derived from MEA electrophysiology, immuno-SERS, and electrochemical sensing using PCA-based dimensionality reduction. Multimodal integration yielded reproducible separation between disease-prone and non-prone assembloids in reduced dimensional space, consistently outperforming single-modality analyses (**Fig. 4a, b**). This separation was maintained across independent assembloid groups, as confirmed by group-wise centroid analysis, supporting the robustness of the integrated feature space (**Fig. 4c**). Examination of principal component loadings revealed that PC1 reflected combined contributions from electrophysiological activity, bioenergetic state, and molecular pathology, whereas PC2 and PC3 captured complementary variations across modalities (**Fig. 4d–f**). By integrating these PCA-weighted features, we derived a composite disease-prone risk score that quantitatively summarized multidimensional vulnerability at the individual assembloid level, consistently discriminating between disease-prone and non-prone assembloids across independent batches (**Fig. 4g**). These results demonstrate that no single analytical dimension is sufficient to capture the full scope of disease-prone divergence — it is the cross-modal integration of transcriptional, molecular, bioenergetic, and electrophysiological dimensions that reveals how genetic and cellular programs translate into functional circuit dysfunction in disease-prone assembloids.

**Fig. 4.**
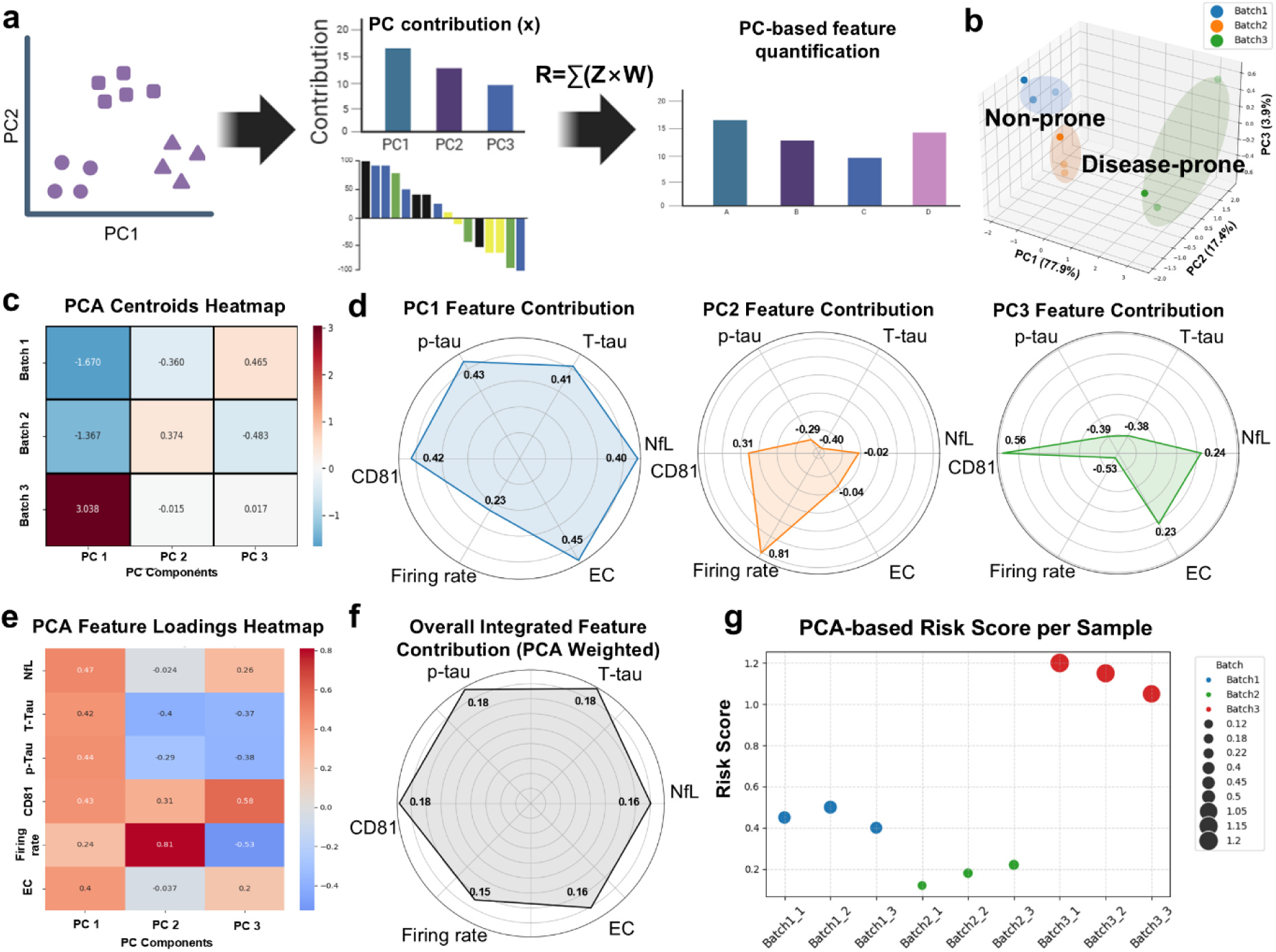
Integrated multimodal analysis identifies disease-prone risk signatures in assembloids. **(a)** Schematic illustration of the PCA-based feature integration workflow. **(b)** PCA of the disease-prone and non-prone groups. **(c)** Group-wise centroid analysis heatmap illustrating reproducible separation across independent assembloid groups. T-tau, Total tau; p-tau, phosphorylated tau. **(d)** Radar plots showing the feature contributions to each principal component (PC1–PC3). **(e)** Heatmap of PCA feature loadings across PC1–PC3, revealing differential contributions from electrophysiological, bioenergetic, and molecular modalities. **(f)** Overall integrated feature contribution across all principal components. **(g)** PCA-based risk score calculated for each assembloid using weighted component contributions.

### Spatial and single-cell transcriptomic characterization of MCAs reveals junction-specific neuronal hyperactivation and neuron-glial uncoupling

To investigate the cellular and molecular basis underlying the disease-prone signatures established by multimodal profiling, we performed spatially resolved laser-activated cell sorting (SLACS) to isolate cells from the forebrain, midbrain, and junction regions of non-prone and disease-prone MCAs, enabling region-specific transcriptomic analysis of each spatial compartment **(Fig. 5a and Extended Data fig. 7-9)**. Gene ontology analysis of junction-upregulated differentially expressed genes (DEGs) revealed significant enrichment of gene ontology terms associated with neuronal connectivity and synaptic function, including neuron projection and glutamatergic synapse **(Fig. 5b)**. Region-specific gene ontology analyses further demonstrated distinct functional and disease-associated signatures across spatial regions **(Extended Data fig. 10)**. We next profiled spatial gene expression at the transcriptomic level across the MCA. Among the three regions, the junction exhibited the largest number of DEGs and showed the greatest DEGs overlap with the forebrain (**Fig. 5c**). Importantly, the junction displayed increased expression of the glutamatergic neuron-associated gene GRIA4, the GABAergic neuron-associated gene GABRG2, and a dopamine signaling-associated gene PDE10A which is highly expressed in GABAergic neurons responsive to dopaminergic input^23–26^. These findings suggest that disease-prone MCAs exhibit a transcriptional program associated with long-range neuronal projection.

**Fig. 5.**
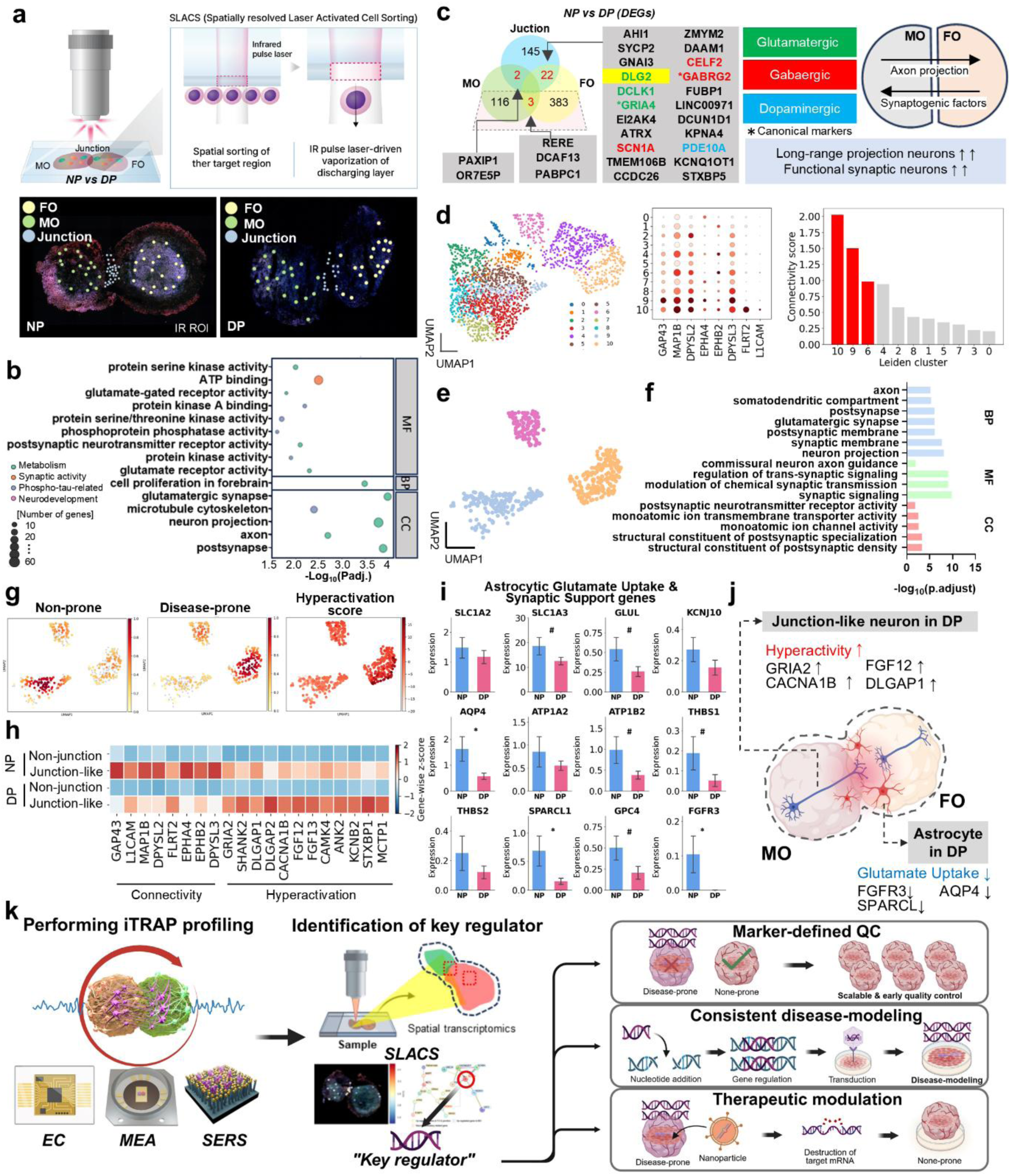
Spatial and single-cell transcriptomic characterization of MCAs reveals junction-specific neuronal hyperactivation and neuron-glial uncoupling. (**a**) Schematic illustration of the SLACS workflow for spatial isolation of assembloids from non-prone and disease-prone assembloids. (**b**) Gene Ontology enrichment analysis of upregulated DEGs in junction region. (**c**) Upregulated DEGs and overlapped DEGs across regions across regions in assembloids. (**d**) Identification of junction-like clusters among neuronal subtypes. (**e**) UMAP of junction-like neuronal cluster. (**f**) Gene Ontology enrichment analysis of upregulated DEGs in junction-like neuronal cluster. MF; molecular function, BP; biological process, CC; cellular component. (**g**) UMAP embedding density and hyperactivation score of junction-like neurons. Benjamini–Hochberg adjusted p-values (FDR) were used. (**h**) Expression heatmap of connectivity and hyperactivation genes (**i**) Expression of astrocytic glutamate uptake and synaptic support genes. *\* P* < 0.05; *# P* <0.1; two-sided p-values; t-test with overestimated variance. **(j)** Schematic summary of junction-like neuronal hyperactivation and impaired astrocytic glutamate uptake identified by single-cell transcriptomic analyses. **(k)** Schematic illustration of the integrated multimodal framework for prospective identification, quality control, and therapeutic application in disease-prone assembloids.

To define the cellular basis of the junction-specific transcriptional program, we analyzed neuronal subtypes using single-cell RNA sequencing. A distinct neuronal cluster enriched for connectivity-associated genes was identified. The three neuronal clusters with the highest connectivity scores were subsequently defined as junction-like neuronal clusters **(Fig. 5d, e)**. Gene ontology analysis revealed enrichment of pathways associated with neuronal projection and synaptic signaling (**Fig. 5f**), indicating enhanced long-range neuronal connectivity and functional synaptic activity within the disease-prone junction. Projection of the hyperactivation signature onto the junction-like neuronal UMAP revealed that disease-prone junction-like neurons occupied regions with higher hyperactivation scores than non-prone cells **(Fig. 5g)**. Disease-prone junction-like neurons exhibited increased expression of hyperactivity-associated genes, including GRIA2, SHANK2, DLGAP1, and CACNA1B, compared with non-prone controls **(Fig. 5h)**. These genes regulate excitatory synaptic transmission and neuronal excitability, consistent with the hyperactive transcriptional state observed in disease-prone assembloids **(Fig. 2c–e)**. Given the hyperactive state of disease-prone junction-like neurons, we next examined astrocytic glutamate uptake and synaptic support gene expression. Disease-prone astrocytes exhibited significantly reduced expression of AQP4, SPARCL1, and FGFR3, with a broader trend toward reduced astrocytic glutamate uptake and synaptic support gene expression compared with non-prone controls (**Fig. 5i**), indicating impaired astrocytic glutamate clearance and synaptic support in disease-prone assembloids.

Collectively, these results support a model in which disease-prone assembloids develop a hyperexcitable junction-like neuronal network accompanied by impaired astrocytic glutamate clearance, ultimately disrupting neuron–astrocyte homeostasis at the forebrain–midbrain interface **(Fig. 5j)**. Together, the convergence of functional, molecular, and spatially resolved transcriptomic evidence positions the corticodopaminergic junction as a candidate region-specific origin of disease-prone divergence, and establishes an integrated multimodal framework for prospective assembloid classification, marker-defined quality control, and precision therapeutic targeting (**Fig. 5k**).

## Discussion

The transition from a healthy to a disease-prone neural circuit state can occur without any detectable morphological change, a principle we demonstrate here in human iPSC-derived brain assembloids, where functional and molecular divergence emerges spontaneously under identical culture conditions. This finding reframes inter-assembloid variability from a technical confound into a biologically meaningful readout of differential neurodegeneration susceptibility, with direct implications for how brain assembloids are characterized and interpreted. Despite low coefficients of variation in circularity and diameter across nine independent batches, multimodal profiling revealed substantial divergence in electrophysiological activity and protein pathology, underscoring that morphological quality control alone is insufficient for characterizing assembloid phenotypes. Critically, disease-prone MCAs exhibited a temporally ordered molecular progression in which NfL elevation preceded tau dysregulation, mirroring the sequential biomarker trajectories reported in pre-symptomatic neurodegeneration in vivo^27^, and suggesting that disease-prone assembloids recapitulate a pre-symptomatic disease state rather than end-stage pathology^21,28^. This early NfL signal, coinciding with localized cortical hyperexcitability in the FO region but preceding global network disruption, suggests that integrated multimodal profiling can capture the earliest detectable features of neurodegeneration, a pre-symptomatic window that remains largely inaccessible in current clinical or animal models and that may prove critical for identifying intervention points before irreversible pathology is established^28^.

Moreover, disease-prone MCAs exhibited systematic decoupling between spontaneous firing rate and tau marker, a disruption that is particularly informative. In non-prone MCAs, spontaneous firing rate was positively correlated with total tau and phosphorylated tau levels, suggesting that tau secretion reflects a physiological correlate of normal neural activity^29,30^, whereas in disease-prone MCAs this relationship is lost, indicating that the link between neural activity and protein homeostasis is fundamentally disrupted. This decoupling would remain entirely undetected by single-modality assessment, underscoring the necessity of integrated analysis. The subsequent elevation of CD81 as a late-stage integrative biomarker, positively correlating with both electrophysiological and molecular markers at 48 hours, further implicates extracellular vesicle dynamics as a downstream component of disease-prone progression^31,32^. At the network level, disease-prone MCAs displayed elevated theta, alpha, beta, and gamma band power alongside a significantly increased theta/delta ratio, recapitulating pathological EEG signatures reported in Alzheimer’s disease and schizophrenia^20^. Preservation of the beta/gamma ratio indicates that oscillatory disruption is predominantly driven by low-to-mid frequency dysregulation, consistent with early-stage rather than end-stage cortical dysfunction. Although MEA recordings do not directly correspond to EEG-defined brainwaves, these brainwave-like oscillatory features suggest that assembloid electrophysiology captures clinically relevant aspects of frequency-dependent network synchronization. Integrated into a PCA-based composite risk score, these multiband features provide a scalable and interpretable metric for phenotype classification that generalizes across independent assembloid batches.

The convergence of electrophysiological hyperexcitability, temporally ordered molecular pathology, bioenergetic dysregulation, and region-specific transcriptional reprogramming collectively defines the disease-prone state, a conclusion supported by the reproducible separation achieved only through multimodal PCA-based integration across these complementary dimensions. Electrochemical sensing of ATP release further demonstrated that bioenergetic dysregulation accompanies the molecular and electrophysiological signatures of disease-prone MCAs, suggesting that metabolic stress may represent an early upstream component of the disease-prone cascade. This integrated framework enables prospective identification of disease-prone assembloids before overt pathology emerges, with direct practical implications. Assembloids can be stratified before experimental use, either excluded to improve reproducibility or enriched to facilitate disease-specific modeling. The identified transcriptional and molecular regulators of disease-prone states further represent actionable targets for precision therapeutic intervention^5,33,34^. More broadly, these findings argue that morphological assessment alone is insufficient as a quality standard for brain assembloid research, and that functional and molecular profiling should be integrated into standard assembloid characterization pipelines.

Building on the multimodal classification framework, spatial transcriptomic analysis provided the mechanistic underpinning of disease-prone divergence. Junction-upregulated DEGs were significantly enriched for GO terms related to neuronal connectivity and synaptic function, including neuron projection and glutamatergic synapse. Additionally, DEGs overlapping between the junction and forebrain were enriched for genes associated with glutamatergic, GABAergic, and dopaminergic signaling, suggesting activation of molecular programs related to midbrain-to-forebrain long-range projections consistent with the developmental trajectory of the mesocortical circuit^35,36^. This transcriptional hyperactivation coincided with impaired astrocytic glutamate clearance at the corticodopaminergic junction, defining a spatially confined neuron-glial uncoupling as a candidate early, region-specific origin of the disease-prone state^37,38^ that could not have been identified in single organoid systems lacking an interregional interface.

Spatial transcriptomic profiling further revealed that forebrain regions were enriched for Alzheimer’s disease and schizophrenia pathways, midbrain-like regions for Parkinson’s disease and epilepsy, and junctional zones for the convergence of multiple disease signatures, reinforcing the translational relevance of the FO–MO assembloid for modeling corticodopaminergic circuit disorders^39,40^. Cell-type-specific transcriptional analysis further revealed that glutamatergic, GABAergic, and dopaminergic neurons each displayed distinct excitability-related gene upregulation in disease-prone assembloids, suggesting that disease-prone divergence involves transcriptional dysregulation across multiple neuronal subtypes and underscoring the importance of single-cell resolution for comprehensively characterizing disease-prone states.

Several limitations warrant consideration. The current analyses were performed using a limited number of assembloid batches and iPSC lines, and the generalizability of disease-prone signatures across diverse genetic backgrounds remains to be established. Furthermore, while the junction-like neuronal population identified here represents a candidate early origin of the disease-prone state, direct causal evidence linking this population to downstream pathology will require targeted perturbation experiments. The current analytical approach is inherently scalable and well-suited for extension to additional iPSC lines, donor backgrounds, and disease-relevant genetic contexts, enabling systematic evaluation of genotype-specific vulnerability signatures.

The present study addresses a fundamental gap in brain assembloid research by demonstrating that functional and molecular heterogeneity encodes biologically meaningful variation in disease susceptibility rather than representing a technical confound to be minimized. This reframing carries broad implications, providing a rationale for prospective phenotypic stratification of assembloids, establishing a multimodal framework that captures pre-symptomatic disease signatures inaccessible to single-modality approaches, and identifying the corticodopaminergic junction as a spatially defined site of early neuron-glial vulnerability. Together, these contributions advance brain assembloid models beyond structural recapitulation toward functional and molecular disease modeling, with direct relevance to a broad spectrum of corticodopaminergic circuit disorders.

In conclusion, this work establishes that phenotypic heterogeneity in brain assembloids is not noise to be minimized but a biologically informative signal reflecting differential susceptibility to neurodegeneration. These findings reposition brain assembloids as windows into pre-symptomatic disease, identify region-specific neuron-glial uncoupling at the corticodopaminergic junction as a candidate early origin of neurodegeneration susceptibility, and open new avenues for mechanistic discovery, inter-individual risk stratification, and precision therapeutic intervention in human-relevant neural models.

## Methods

### Generation of dorso forebrain organoids

Dorso forebrain organoids (FOs) were generated from human induced pluripotent stem cells (iPSCs) (BIONi010-C) as previously described^5,41^. An alpha-tubulin-GFP tagged iPSC line (Coriell Institute, AICS-0012) was used to generate fluorescent organoids. The hiPSCs were initially maintained on Matrigel hESC-qualified Matrix-coated plates (SPL, 20100) in mTeSR Plus (StemCell Technologies, ST100-0276). To create embryoid bodies (EBs), hiPSCs were detached using ReLeSR, and the resulting colonies were dissociated into single cells in AggreWell EB Formation Medium (Stemcell Technologies, ST05893) supplemented with the ROCK inhibitor Y-27632 (Stemcell Technologies, ST72304). On Day 0, 1.5 × 10^6^ cells were seeded into each well of an AggreWell800 24-well plate (StemCell Technologies, 34811). The following day, the medium was replaced with EB Formation Medium only. From Day 2 to Day 5, daily medium changes were performed using DMEM/F-12 with GlutaMAX (Gibco, 10565-018) supplemented with 20% KnockOut Serum Replacement (Gibco, A3181501), 1% MEM Non-Essential Amino Acids Solution (Gibco, 11140050), 0.1 mM 2-mercaptoethanol (Gibco, 21985023), and 100 U/mL penicillin and 100 μg/mL streptomycin (Merck, P4333), as well as SMAD inhibitors, dorsomorphin (10 μM; Merck, P5499), and SB-431542 (10 μM; TOCRIS, 1614). On Day 6, EBs were transferred to 100-mm dishes containing Neurobasal-A Medium (Gibco, 10888-022), B-27 Supplement minus vitamin A (Gibco, 12587010), 100 U/mL penicillin, 100 μg/mL streptomycin, GlutaMAX (Gibco, 35050-061), and 0.5% (v/v) Matrigel Basement Membrane Matrix (Corning, 354234), and cultured on an orbital shaker. The following day, individual EBs were transferred to separate wells of a 96-well ultra-low attachment microplate (Corning, 7007). From Day 6 to Day 15, the medium was changed daily, supplemented with 20 ng/ml Epidermal Growth Factor (EGF; Peprotech, AF-100-15-500 μg) and 20 ng/ml Fibroblast Growth Factor basic (bFGF; R&D Systems 100-18B). Between Day 16 and Day 24, the medium was refreshed every other day, with the same growth-factor composition. Starting on Day 25, EGF and bFGF were withdrawn and substituted with 20 ng/mL Brain-Derived Neurotrophic Factor (BDNF; Peprotech, 450-02) and 20 ng/mL Neurotrophin-3 (NT-3; Peprotech, 450-03), with medium changes continuing on an alternate-day schedule through Day 42. Subsequently (from Day 43 onward), the organoids were maintained in a growth-factor-free neural medium, refreshed every 4 days.

### Generation of midbrain-like organoids

Midbrain-like organoids (MOs) were generated using a modified version of a previously described protocol^42^, utilizing ApoE3/4 iPSCs and RFP-tagged alpha-tubulin line (Coriell Institute, AICS-0031-035). EBs were initially formed using the forebrain organoid procedure and subsequently plated into ultra-low attachment 96-well plates (Corning, 7007) containing 50 μL of neural induction medium (IM). IM consists of DMEM/F12 (Thermo Fisher Scientific, 2645233) and Neurobasal medium (Thermo Fisher Scientific, 2661481) in a 1:1 ratio, supplemented with N2 (17502048), B27 without vitamin A (2596532), GlutaMAX (35050061, Gibco), MEM non-essential amino acids (11140, Thermo Fisher Scientific), and β-mercaptoethanol (21985023, Gibco). Additional components included heparin (Sigma-Aldrich, H3149), SB431542 (Tocris, 1614), Noggin (Prospec, CYT-475), CHIR99021 (Tocris, 4423), Y27632 (Calbiochem), and growth factor-reduced (GFR) Matrigel. On Day 6, an additional 100 μL of IM supplemented with SHH-C25II (R&D Systems, 464-SH) and FGF8 (R&D Systems, 423-F8) was added to each well to induce neuronal patterning. On Day 9, the medium was replaced with tissue growth (TG+) medium comprising Neurobasal medium with N2, B27 without vitamin A, GlutaMAX, MEM non-essential amino acids, and β-mercaptoethanol, further supplemented with insulin, laminin, SHH-C25II, FGF8, and GFR Matrigel, and incubated for 24 h. On Day 10, the organoids were transferred to ultra-low attachment 24-well plates (Corning, 3473), with each well containing 500 μL of organoid growth (OG+) medium comprising Neurobasal medium with N2, B27 without vitamin A, GlutaMAX, MEM NEAA, and β-mercaptoethanol, further supplemented with BDNF, GDNF, ascorbic acid, dibutyryl cAMP (db-cAMP), and GFR Matrigel. The medium was then replaced every three days.

### Generation of dorso forebrain-midbrain mesocortical assembloids

Day 100 FOs and Day 100 MOs were physically fused to generate dorso forebrain-midbrain mesocortical assembloids (MCAs) under standard culture conditions (37°C, 5% CO₂). To facilitate initial cell–cell interactions, FOs and MOs were placed together in individual wells of ultra-low attachment 24-well plates (Corning, 3473), with the plates maintained at a slight tilt during incubation to promote gravitational contact at the organoid interface. After 1–2 days under static conditions, once cell adhesion was established and fusion initiated, the plates were transferred to an orbital shaker at 90 rpm to enhance nutrient and oxygen exchange and promote structural maturation. To support cellular reorganization at the FO–MO junction interface and enable neurite extension for functional integration across regions, the assembloids were cultured for two weeks prior to downstream multimodal analysis.

### Fabrication of SERS substrates

Gold-coated zinc oxide (ZnO) nanowire substrates were fabricated for SERS analysis^41^. Silicon wafers (2.5 × 2.5 mm²) were diced and sequentially cleaned with acetone, isopropyl alcohol, and distilled water (DW). The cleaned Si wafers were seeded by spin-coating with 25 mM zinc acetate dihydrate (Zn(CH₃COO)₂·2H₂O, 96459, Sigma Aldrich) in isopropyl alcohol. ZnO nanowires were grown on the seeded substrates through hydrothermal synthesis at 90°C for 1 h, using 40 mM zinc nitrate hexahydrate (Zn(NO₃)₂·6H₂O, 228737, Sigma Aldrich) and 25 mM hexamethylenetetramine (HMTA, 33233, Sigma Aldrich) as the reactant and stabilizer, respectively. After synthesis, the substrates were rinsed twice with DI water, dried under nitrogen flow, and coated with a 100-nm gold layer via thermal evaporation.

### Antibody functionalization for SERS

SERS substrates were functionalized with half-antibody fragments to enhance binding specificity. Individual antibodies–anti-CD81, anti-NfL, and anti-tau (MN1000, Thermo Fisher Scientific)–were diluted to 0.05 mg/mL in phosphate-buffered saline (PBS, P3813, Sigma Aldrich). Tris (2-carboxyethyl)phosphine (0.02 mM; Bond-Breaker TCEP Solution, 77720, Thermo Fisher Scientific) was added at a 1:1 volume ratio and incubated for 1 h at room temperature to cleave the disulfide bonds in the antibody hinge region, generating half-antibody fragments. 10 µL of each half-antibody solution was applied to individual SERS substrates and incubated overnight at 4°C. The functionalized substrates were then rinsed with DI water, dried under nitrogen, blocked with a 1× Pierce clear milk blocking solution (37587; Thermo Fisher Scientific) for 4 h at RT, and rinsed again before use.

### SERS detection

The immuno-SERS substrates co-localized on the HD-MEA chip were retrieved at 6, 24, and 48 h, rinsed twice with DI water, and dried under nitrogen flow. SERS spectra were acquired using a Raman spectroscope (NS 200, Nanoscope Systems) with 785 nm laser excitation (1.8 mW) through a 50× objective lens (NA 0.8). Each spectrum was acquired for 1 s. Baseline correction was performed using the asymmetric least-squares method, and the corrected spectra were smoothed with a Savitzky-Golay filter. For disease-prone classification, normalized SERS intensities were used for marker quantification. Threshold values for each marker were determined based on the distribution of normalized intensities across all assembloids, with cutoffs set at the upper quartile of the overall population to maximize separation between groups. Assembloids exhibiting normalized intensities exceeding the defined threshold for at least two markers (T-tau > 1.0, p-tau > 0.4, or NfL > 0.5) were designated disease-prone, whereas those maintaining normalized intensities below all thresholds were designated non-prone.

### MEA preparation and electrophysiological recordings

MaxOne MEAs (MaxWell Biosystems) were sterilized by immersion in 70% ethanol for 30 min, followed by three rinses with sterile DW. To enhance tissue adhesion, the arrays were coated with 0.05% (v/v) polyethylenimine (Sigma-Aldrich, #P3143) in borate buffer (pH 8.5; Thermo Scientific, #28341) for 40 min at RT, rinsed with DW, and air-dried. Laminin (#L6274; Sigma-Aldrich) was applied to the central recording area. The neural activity of the assembloids was detected using MaxOne-MEA through sequential high-density activity scans covering all array electrodes. Recordings were acquired at a sampling rate of 10 kHz with a lower cutoff frequency of approximately 1 Hz. Recording sites were selected based on multiunit activity and quantified for each electrode using a sliding-window spike detection algorithm with a threshold set at five times the root mean square noise of the bandpass-filtered signal^19^.

### Electrode data acquisition

Raw voltage traces were extracted from MEA recordings using the Maxwell Biosystems proprietary MATLAB toolbox. Data were sampled at the native acquisition frequency of the system (20 kHz), and continuous segments were retrieved from the raw datasets. In addition to unprocessed traces, bandpass-filtered signals (300–3000 Hz) were obtained for spike detection and electrode selection^43^.

### Frequency-domain analysis

Raw voltage traces were filtered using a zero-phase Butterworth bandpass filter (1–100 Hz) to isolate physiologically relevant oscillatory activity, while removing slow drifts and high-frequency noise. Filtered traces were transformed into the frequency domain using a fast Fourier transform (FFT) in MATLAB. For each electrode, the power spectral density (PSD) was computed as the squared magnitude of the FFT, normalized by the signal length.

### Fabrication of electrochemical sensing analysis

The electrochemical (EC) sensing analysis was fabricated using indium tin oxide (ITO) glass substrates, gold chloride trihydrate (AuCl₃), poly(ethylene glycol) 200 (PEG 200), Triton X-100, DPBS (Sigma-Aldrich), polydimethylsiloxane (PDMS), and plastic chambers sourced from U.I.D, Sigma-Aldrich, and Dow Corning Corp. ITO substrates (1.2 cm × 1.7 cm; thickness: 0.07 cm; electrical resistance: 8 Ω) were sequentially cleaned by ultrasonication in 1% Triton X-100 solution, deionized (DI) water, and 70% ethanol. A plastic chamber was affixed to the ITO electrode using PDMS (10:1 base-to-curing-agent ratio), which served as a biocompatible scaffold for electrochemical gold deposition and organoid culture. The gold deposition solution was prepared by mixing a 5 mM gold (III) chloride solution with PEG 200 at a 50:1 ratio. This solution was electrochemically deposited onto the ITO substrate using a multistep potential (MSP) protocol on an electrochemical workstation (EG Technology) for 120 s, as previously described^44^. After deposition, the chips were rinsed with 70% ethanol and sterilized under ultraviolet light for 40 min. The gold nanostructured surface was then coated with Matrigel (Corning) diluted 1:80 and incubated at 37°C for at least 1 h before organoid placement.

### EC monitoring of brain assembloids

EC monitoring was performed using differential pulse voltammetry (DPV) on a DY2013 Potentiostat (EG Technology). The Matrigel/gold nanostructure/ITO assembly served as the working electrode, with a platinum wire and an Ag/AgCl (3.5 M KCl) electrode as the counter and reference electrodes, respectively. Before measurement, the culture medium was replaced with fresh medium to minimize interference from redox-active metabolites. For precise quantification, the DPV signals were set as follows: initial E (V) = –0.3, final E (V) = 0.5, step E (V) = 0.005, and pulse period (s) = 0.2. All measurements were conducted at room temperature (RT). The calculated Ip values were analyzed by subtracting the baseline current from the current at Ep = –0.25 V.

### Principal component analysis (PCA)-based weighted score calculation

The EEG frequency-band powers (Delta, Theta, Alpha, Beta and Gamma) and measured values (T-tau, p-tau, EC, firing rate, NfL, and CD81) were collected. Feature values were standardized across samples using z-score normalization before downstream analysis. Feature contributions were calculated by combining the PCA loadings with the explained variance of each retained component. For the feature *i*, the signed contribution *C_i_*was defined as follows:

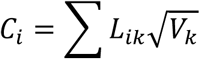

*L_ik_* denotes the loading of the feature *i*on the principal component *k*, and *V_k_* represents the explained variance ratio of the component *k*. The square root of the explained variance ratio was used to weigh the component loadings, ensuring scale consistency with the principal component scores^45^.

### PCA-based risk score calculation

Sample-wise risk scores were computed as a weighted linear combination of normalized EEG features. For sample *j*, the risk score *R_j_* was calculated as follows:

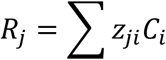

where *Z_ji_* is the z-score-normalized value of feature *i* in sample *j*, and *C_i_* is the PCA-based feature contribution defined above.

### Spatial sequencing analysis

Genes with adjusted *P* < 0.05 were defined as differentially expressed genes (DEGs). The DEGs were subjected to gene ontology (GO) enrichment and protein-protein interaction (PPI) network analyses. GO analysis was performed using the ToppGene database (https://toppgene.cchmc.org/) and PPI analysis was conducted using the STRING database (https://string-db.org/).

### Immunohistochemistry

Organoids were rinsed with PBS, fixed overnight at 4°C in 4% paraformaldehyde (PFA), and rinsed again with PBS. Samples were cryoprotected in 30% sucrose at 4°C for 72 h, embedded in FSC 22 compound (Leica, 3801480) within a cryomold, and frozen. Organoids were then cryosectioned at 40 μm thickness, and sections were rinsed with PBS and permeabilized with 0.3% Triton X-100 (Merck, X100) in PBS for 30 min at RT. For blocking, the sections were incubated in 5% normal horse serum (H0146; Sigma) in PBS for 1 h at RT. Primary antibodies, diluted in blocking buffer, were applied overnight at 4°C. Fluorescently labeled secondary antibodies (1:500 in 3% BSA/PBS) were then applied for 1 h at RT. After washing, the sections were mounted on slides. The following primary antibodies were used: anti-AQP4 (1:500; SYSY, 429 009), anti-PSD95 (1:500; Invitrogen, MA1-045), anti-NeuN (1:500; Cell Signaling Technology, 24307T), and anti-DAT (1:500; Sigma-Aldrich, MAB369).

### Spatial region isolation via SLACS

Spatially defined tissue regions were isolated using CosmoSort (Meteor Biotech, Republic of Korea), a commercially available spatial cell-sorting analysis that utilizes infrared pulse laser-based ejection for the non-destructive retrieval of histologically annotated regions. CosmoSort uses an infrared pulse laser-based ejection mechanism to selectively transfer annotated regions of interest (ROIs) into designated collection wells for downstream RNA analysis. Tissue sections were subjected to immunofluorescence staining, and whole-slide images were processed using Astromapper, a proprietary coordinate mapping software that enables digital overlay and precise ROI localization. Astromapper-generated coordinate files were imported into the CosmoSort system, which automatically aligned physical slides with annotated images, and performed laser-mediated ejections of each ROI into individual collection wells for downstream analyses. A total of 90 ROIs (30 per assembloid × 3 assembloids, 10 from each region) were sequenced for spatial transcriptomic analysis.

### RNA-seq data processing and analysis

RNA sequencing data were processed using a standard paired-end workflow aligned with the human genome (hg38/GRCh38). Adapter trimming and quality control were performed using Cutadapt. The processed reads were aligned to the reference genome using a STAR aligner to generate BAM files. Gene-level read counts were quantified using FeatureCounts, while RSEM was used for isoform-level expression quantification to generate FPKM and TPM values. Differential gene expression analysis was performed using DESeq2 in R.

### Differential gene expression analysis

Raw gene-level read counts from FeatureCounts were used as input for DESeq2 (v1.38.0) in R. Low-quality genes were filtered to improve statistical power; genes with fewer than two reads across samples or mean expression below 0.5 were excluded. Count data were imported using the DESeqDataSetfromMatrix, where type represented the biological condition of interest for each sample. Normalization was performed using the median-of-ratios method to account for sequencing depth and compositional biases. The DESeq2 estimated the size factors and dispersions, and performed negative binomial Wald tests for differential expression. Log₂ fold changes were shrunk using the apeglm and ashr methods to improve the effect size estimation for low-count genes. Adjusted p-values were corrected for multiple testing using the Benjamini–Hochberg false discovery rate (FDR) method. Genes with |log₂(fold change)| ≥ 0.58 (equivalent to 1.5-fold change) and FDR-adjusted *P* < 0.05 were considered significantly differentially expressed.

### Data acquisition and analysis

ATP levels were monitored in real-time via EC readouts, and spontaneous neural activity was recorded simultaneously using the MEA. SERS-derived spectral intensities corresponding to T-tau, p-tau, and NfL were quantified from baseline-corrected spectra and subsequently used for normalized intensity-based disease-prone classification as described above. All data streams–EC, electrophysiological, and molecular–were temporally aligned and processed using custom MATLAB scripts for downstream statistical analyses. Spatial transcriptomic data obtained via SLACS were integrated with functional and molecular readouts to enable comprehensive multimodal characterization of each assembloid.

### Statistical analysis

All quantitative data are presented as mean ± SEM and statistical analyses were performed using unpaired Student’s *t*-test or one-way ANOVA followed by Tukey’s post hoc test, as appropriate. Statistical significance was set at *P* < 0.05.

## Data availability

The spatial sequencing data from this study (related to Fig. 5 and Extended Data Figs. 7–10) have been deposited in the GEO accession number GSE316685. The single-cell RNA sequencing data from this study (related to Fig. 1, Fig. 5, and Extended Data fig. 2) are currently being processed and will be deposited in GEO upon completion. The datasets generated and analyzed are available from the corresponding author upon appropriate request.

## Code availability

N/A

## Acknowledgments

This work was supported by the National Research Foundation of Korea (NRF), funded by the Korean government (MSIT) (grant nos. RS-2022-NR072469, RS-2023-00266110, and RS-2026-25520892 to J.-C.P.). Additional support was provided by the NRF grant funded by MSIT (grant no. RS-2024-00462912). This research was further supported by NRF grants funded by MSIT (grant no. RS-2024-00454407); the Technology Innovation Program and the Industrial Strategic Technology Development Program funded by the Ministry of Trade, Industry and Energy (MOTIE), Republic of Korea (grant nos. 20024391, RS-2024-00451981, and RS-2024-00508420); and the Korea–US Collaborative Research Fund (KUCRF), funded by the Ministry of Science and ICT and the Ministry of Health & Welfare, Republic of Korea (grant no. RS-2024-00468338).

## Author contributions

S.K., R.K., T.L., and J.-C.P. conceived and designed the study. L.P.L., I.K., and J.-C.P. provided the conceptual framework for integrated multimodal monitoring of brain organoids. S.K. coordinated all the experiments. T.L. generated brain assembloids and performed imaging analyses. Y.K. contributed to the SERS analyses. R.K. analyzed the MEA and spatial transcriptomic data, and J.N. assisted with the scRNA-seq analyses. C.D.K., K.M.K., and T.H.K. contributed to the electrophysiological measurements. S.L., D.K., H.O., and A.C.L. performed SLACS and generated the spatial omics data. B.P., L.P.L., I.K., and J.-C.P. provided scientific advice and conceptual input. S.K., R.K., T.L., Y.K., L.P.L., I.K., and J.-C.P. wrote the manuscript with input from all authors. L.P.L., I.K., and J.-C.P. supervised the study and provided overall guidance.

## Extended Data Figures

**Extended Data fig. 1.**
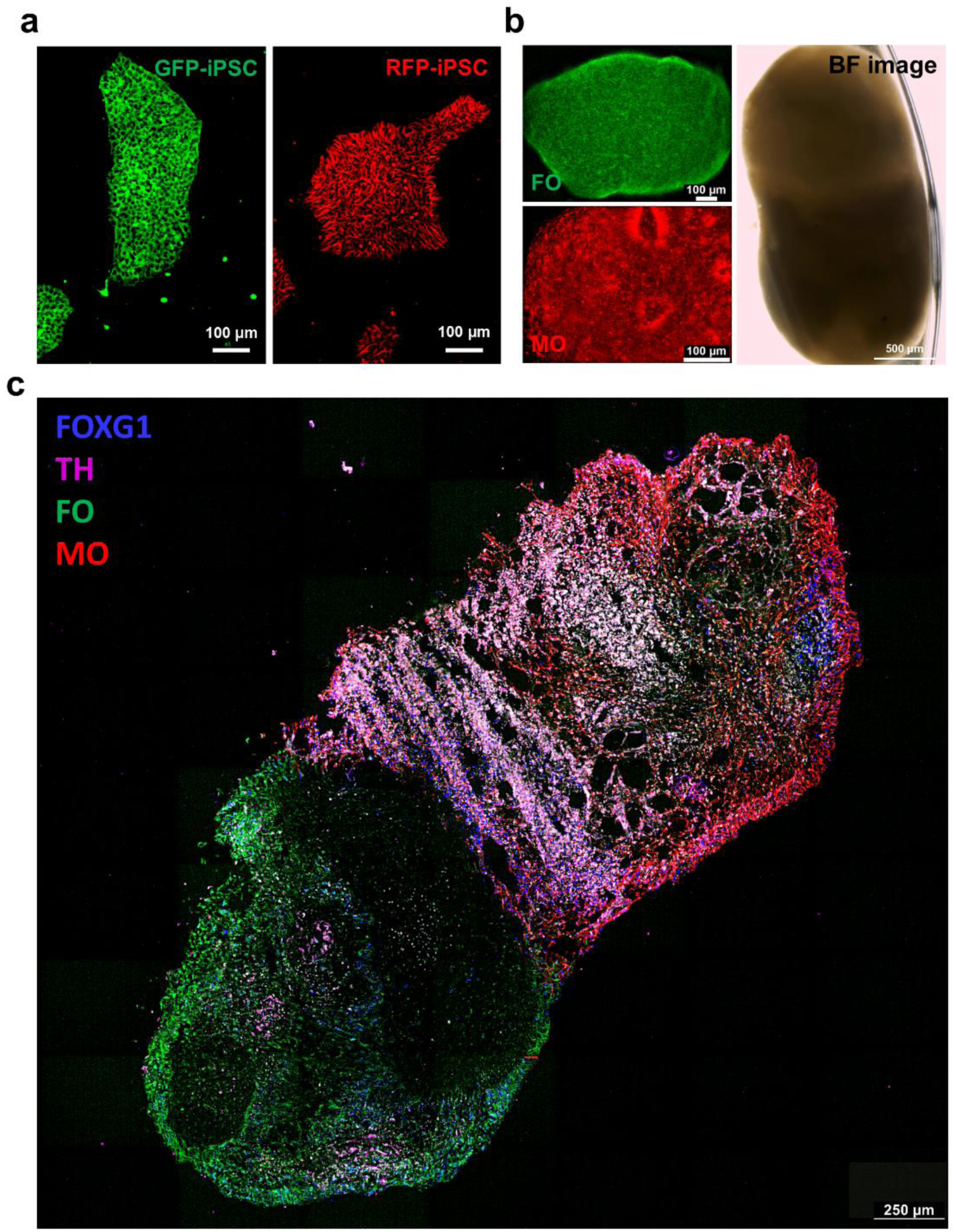
Validation of FO, MO, and assembloids. **(a)** GFP and RFP tagged-tubulin iPSC (Coriell Institute; AICS-0012, AICS-0031-035) were used to visualize neural connectivity. **(b)** GFP-FO and RFP-MO were generated seperately. Brightfield image of daf. 21 brain assembloids using FO and MO. **(c)** Immunohistochemistry was performed to distinguish FO and MO using region-specific markers. FOXG1(abcam, ab18259), a marker of telecephalon and cerebral cortex, was expressed in the distal GFP-FO, whereas TH (abcam, ab6211), a dopaminergic marker, was strongly expressed in the RFP-MO.

**Extended Data fig. 2.**
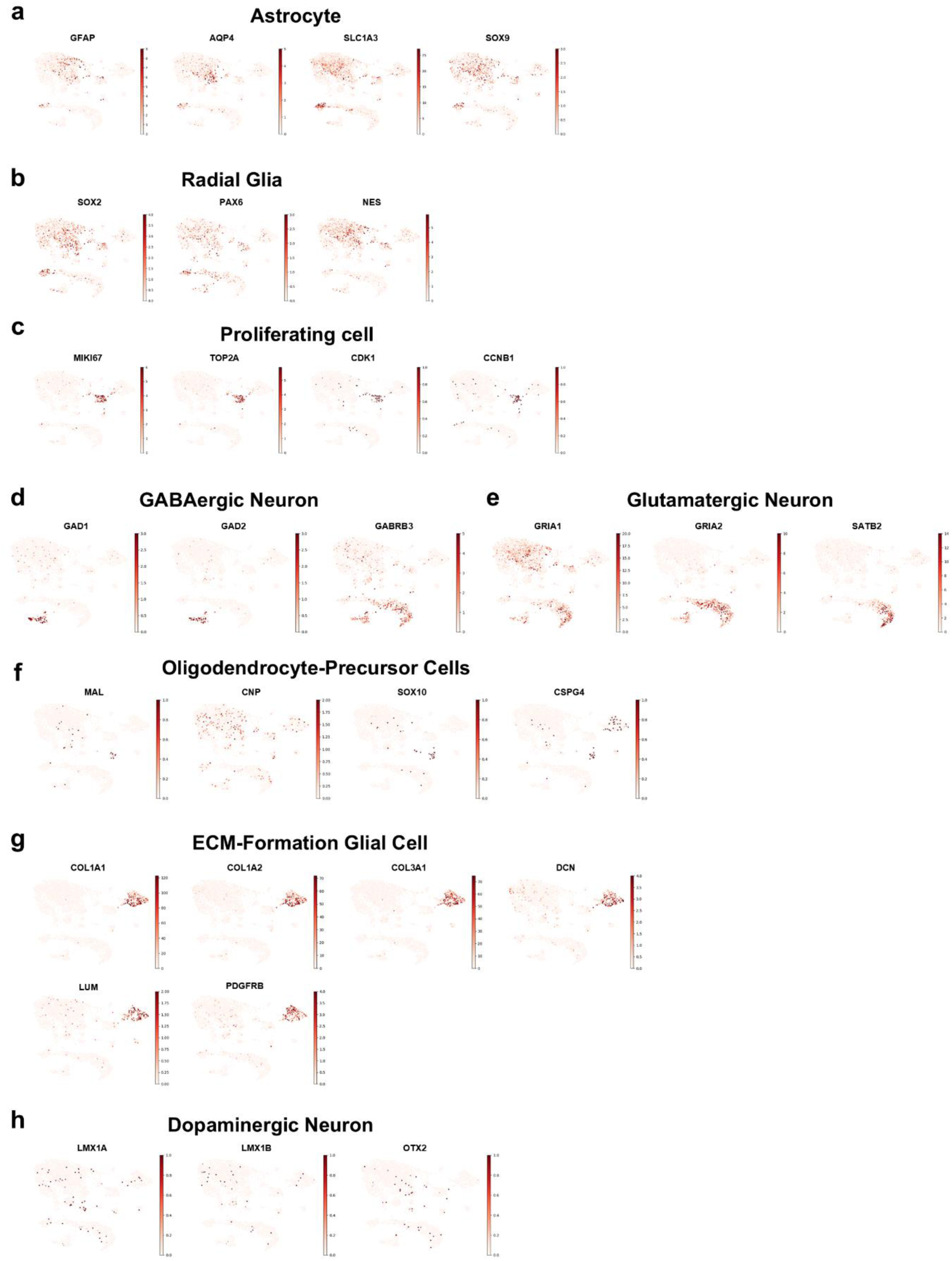
Marker genes used for cell-type annotation visualized on UMAP. **(a-h)** Marker genes expression of each cell types.

**Extended Data fig. 3.**
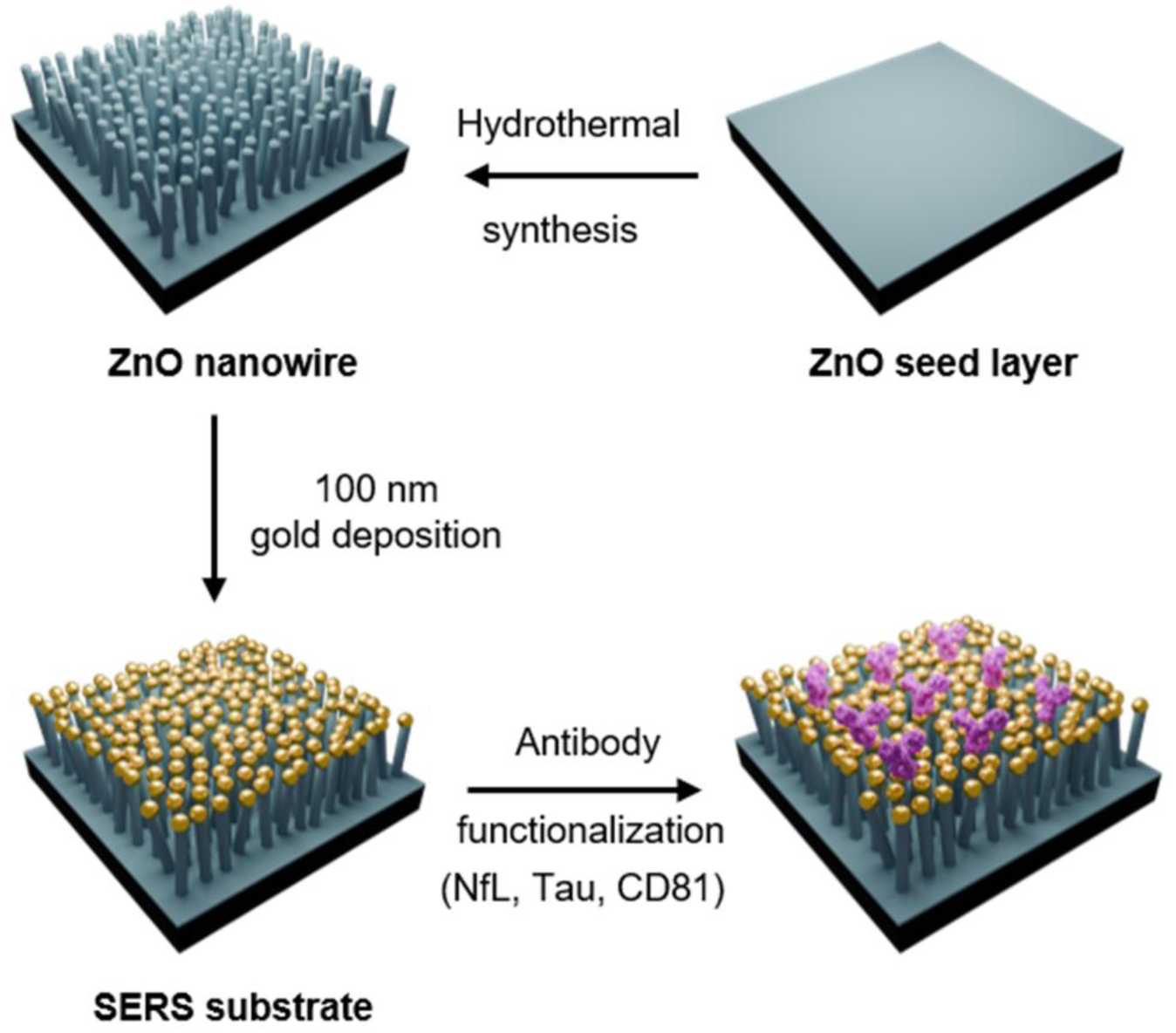
Schematic of SERS substrate fabrication. ZnO nanowires are deposited via hydrothermal synthesis, followed by ZnO seed layer formation. The substrate is then functionalized with antibodies (NfL, Tau, CD81) for selective biomarker detection using MEA-integrated SERS platform.

**Extended Data fig. 4.**
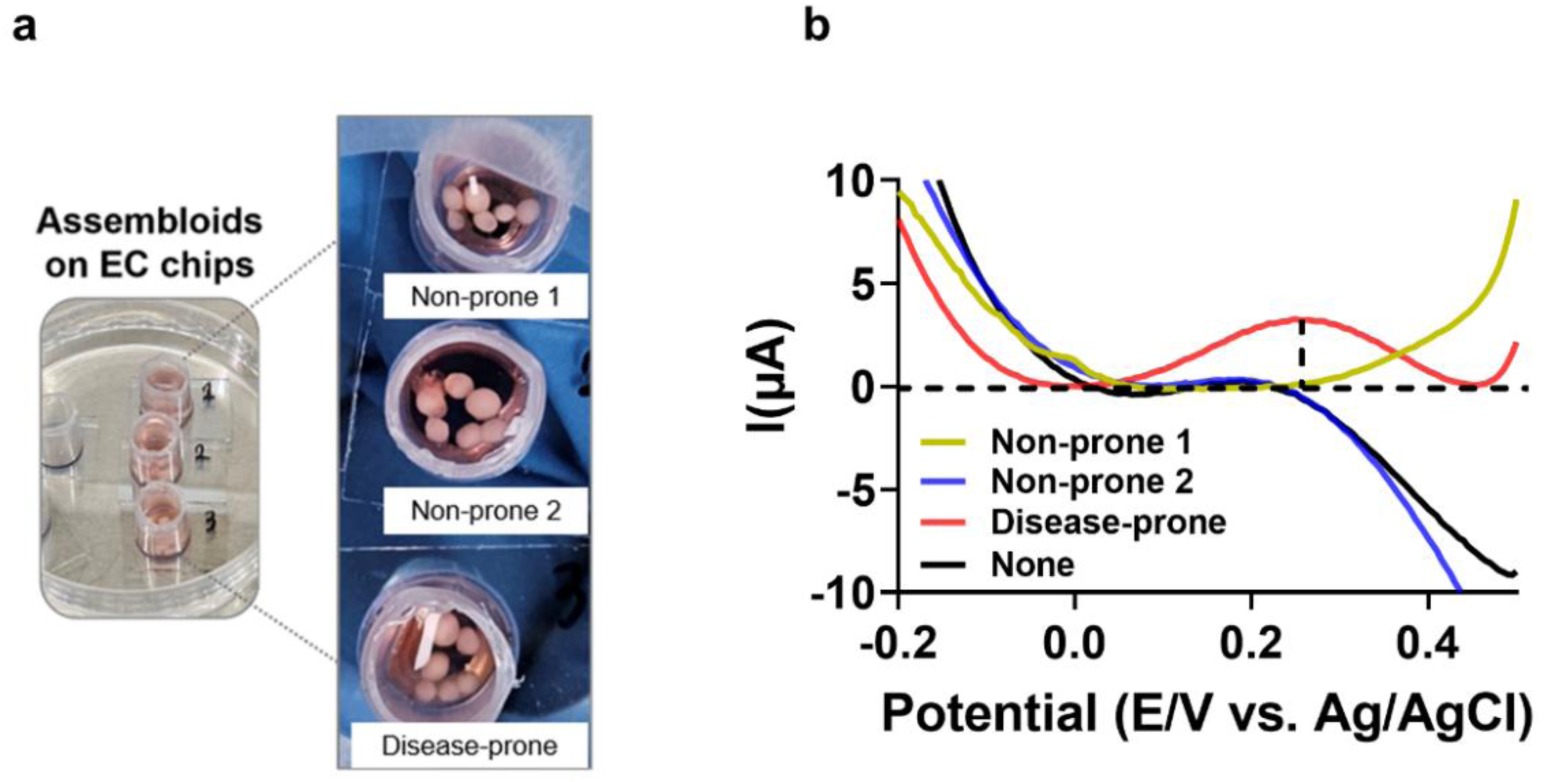
Electrochemical (EC) validation using brain assembloids and organoids. **(a)** Representative images of assembloids cultured on EC chips and the corresponding EC profiles showing group-dependent divergence in ATP release. **(b)** Disease-prone assembloids showing elevated bioenergetic activity compared to non-prone assembloids. Consistent peak positions (∼0.2V vs Ag/AgCl) demonstrate reliable detection across multiple measurements, validating the EC chip for organoid-based disease modeling.

**Extended Data fig. 5.**
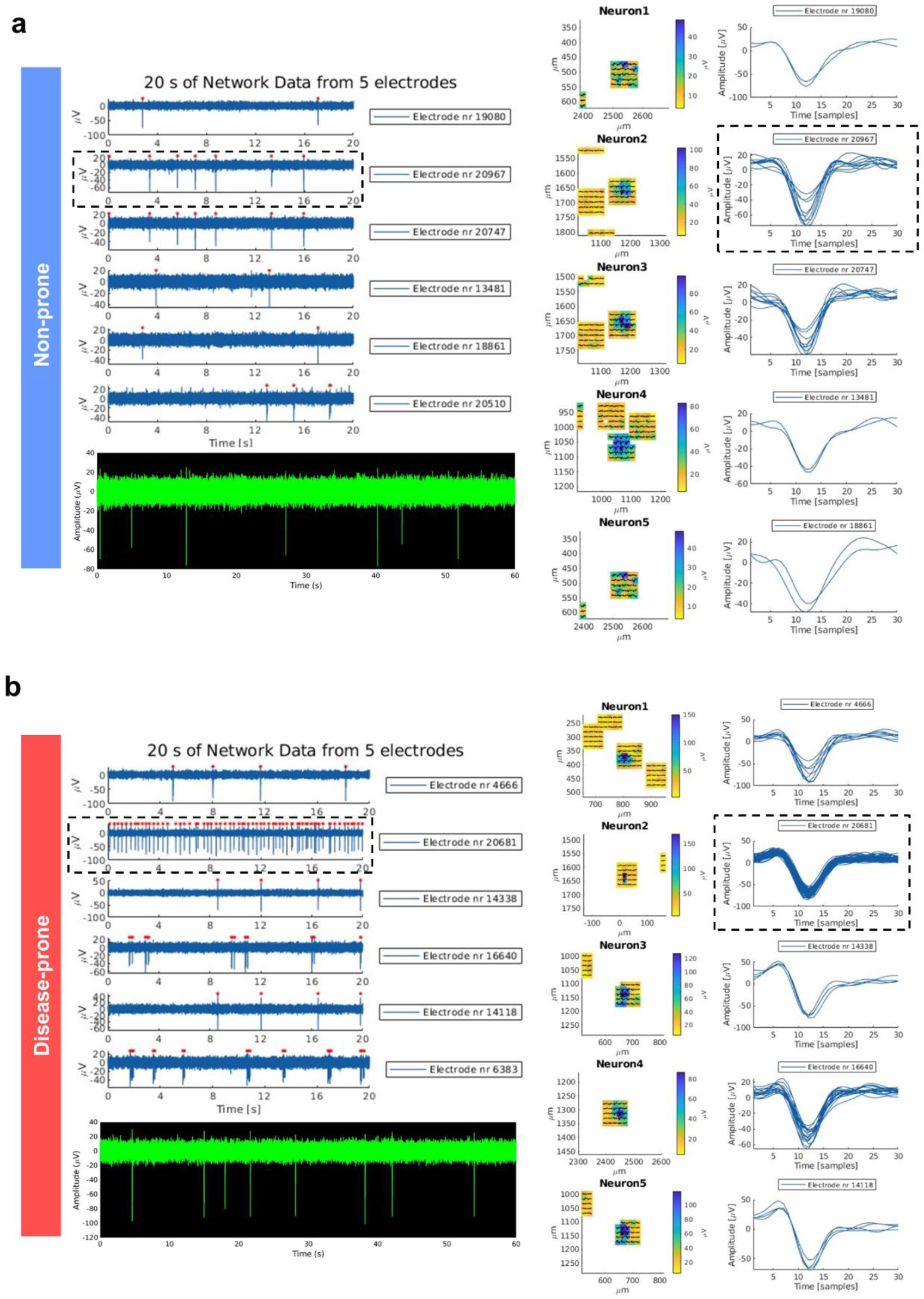
Representative MEA recordings from non-prone and disease-prone assembloids. **(a-b)** Representative network activity traces from multiple electrodes, spike waveforms, and spatial electrode maps recorded from non-prone (top) and disease-prone (bottom) assembloids. Panels outlined with dashed boxes are also shown in Fig. 2c.

**Extended Data fig. 6.**
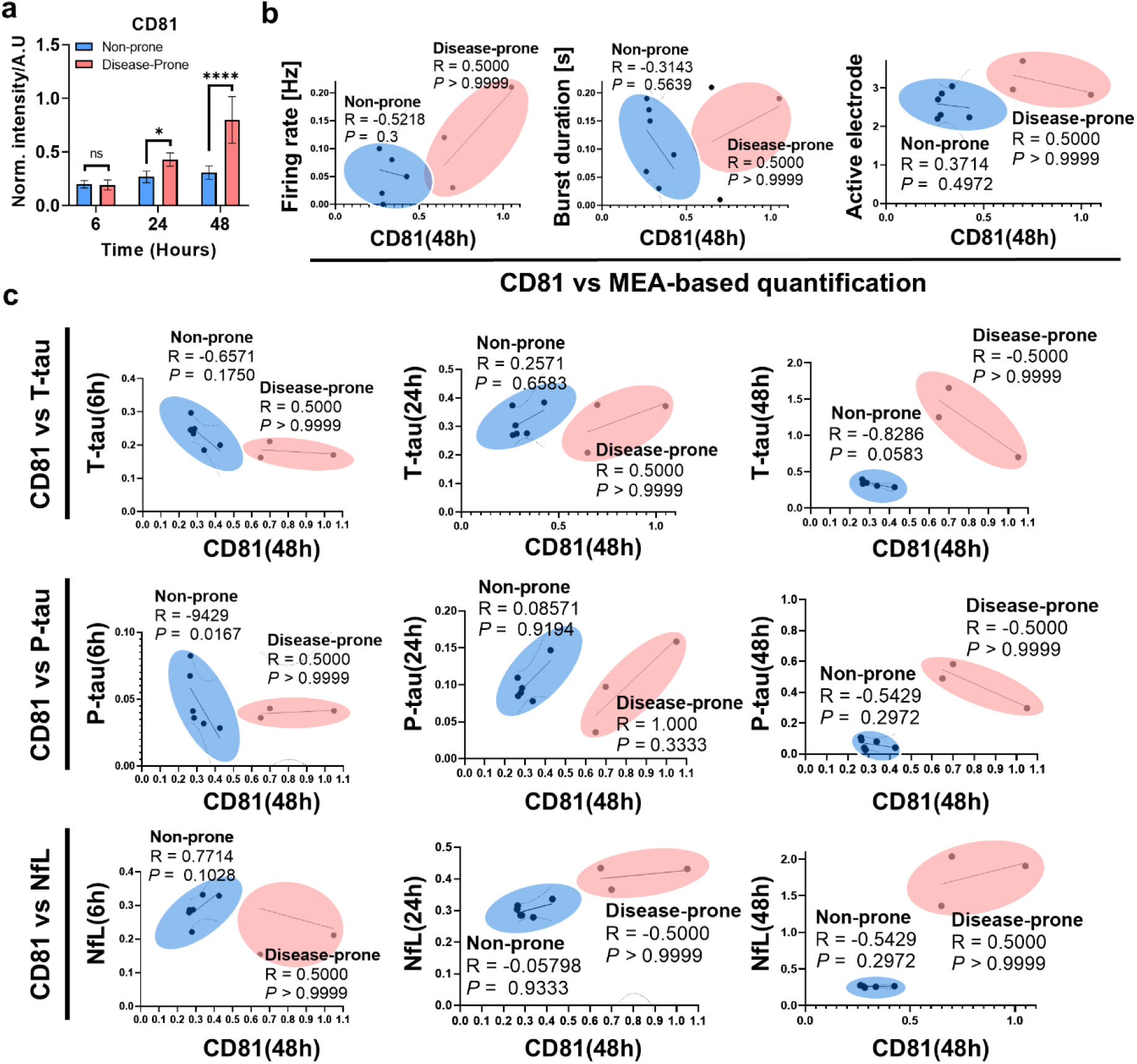
Elevated levels of the exosomal marker CD81 associate with neurodegenerative and functional signatures in disease-prone assembloids. **(a)** CD81, a tetraspanin protein and canonical marker of extracellular vesicles, was profiled via immuno-SERS across all assembloids at 6, 24, and 48 hours. Disease-prone assembloids showed significantly elevated CD81 signals compared to non-prone assembloids. Statistical significance: ns, not significant; * *P* < 0.05; **** *P* < 0.0001. **(b)** Scatterplots showing correlations between CD81 levels (48 hours) and MEA-based functional measurements including burst duration, firing rate, and active electrode count. **(c)** Scatterplots showing Spearman correlations between CD81 levels (48 hours) and immuno-SERS-based neurodegeneration markers (T-tau, p-tau, NfL). Spearman correlation coefficients and p-values are indicated for each comparison. Statistical significance was defined as *P* < 0.05.

**Extended Data fig. 7.**
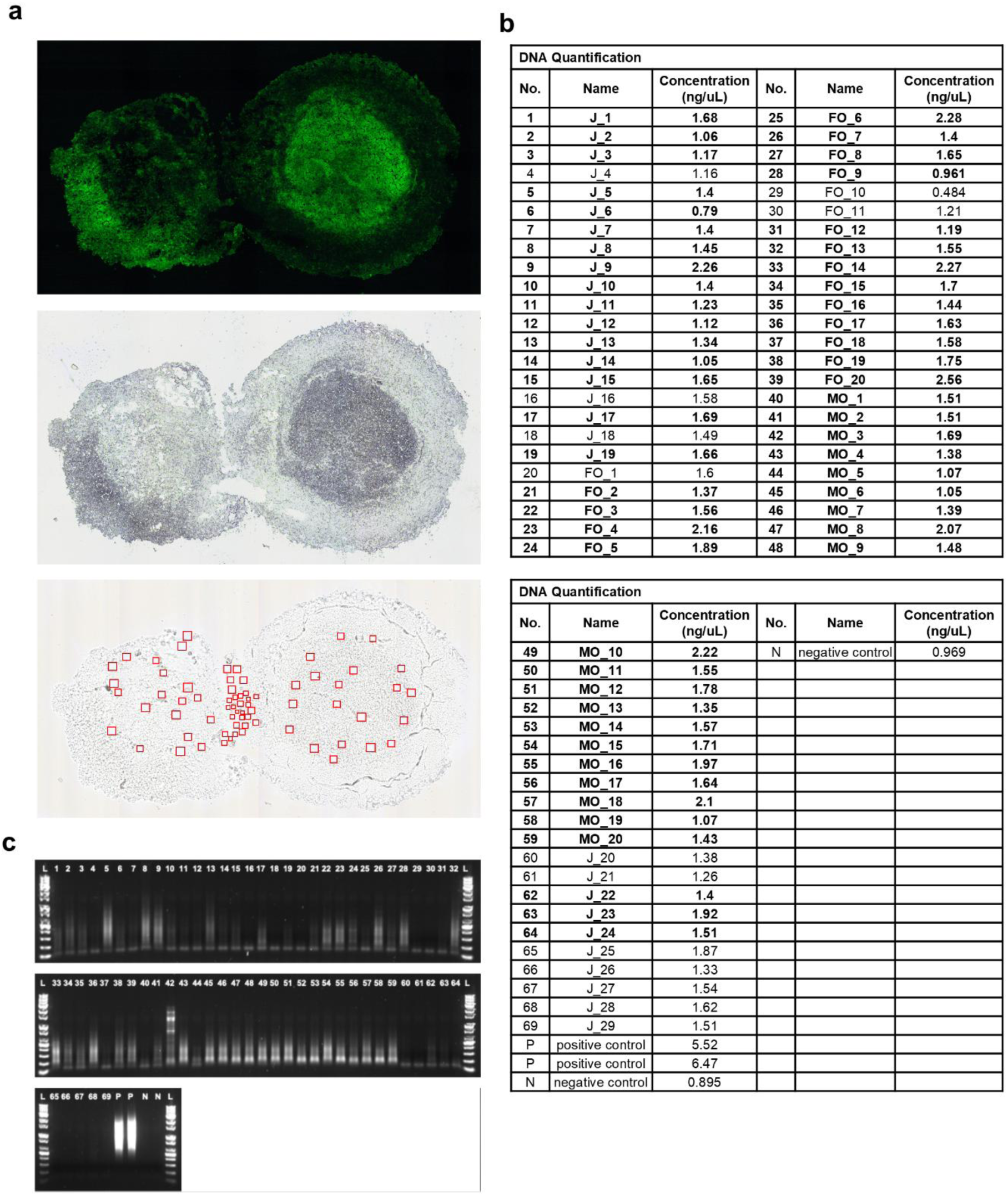
Quality control and sample information for spatially resolved transcriptomic analysis. **(a)** Three regions were analyzed: MO, FO, and junction (J). **(b)** A total of 69 ROIs were isolated (20 MO, 20 FO, and 29 J), of which 51 ROIs (20 MO, 17 FO, and 14 J) passed QC. **(c)** QC pass was determined based on the presence of a smeared band pattern in the electrophoresis profile. Sequencing was performed on 30 ROIs in total (10 from each group). RNA-seq quality assessment results, including RNA concentration and electrophoresis profiles, are shown in the same figure.

**Extended Data fig. 8.**
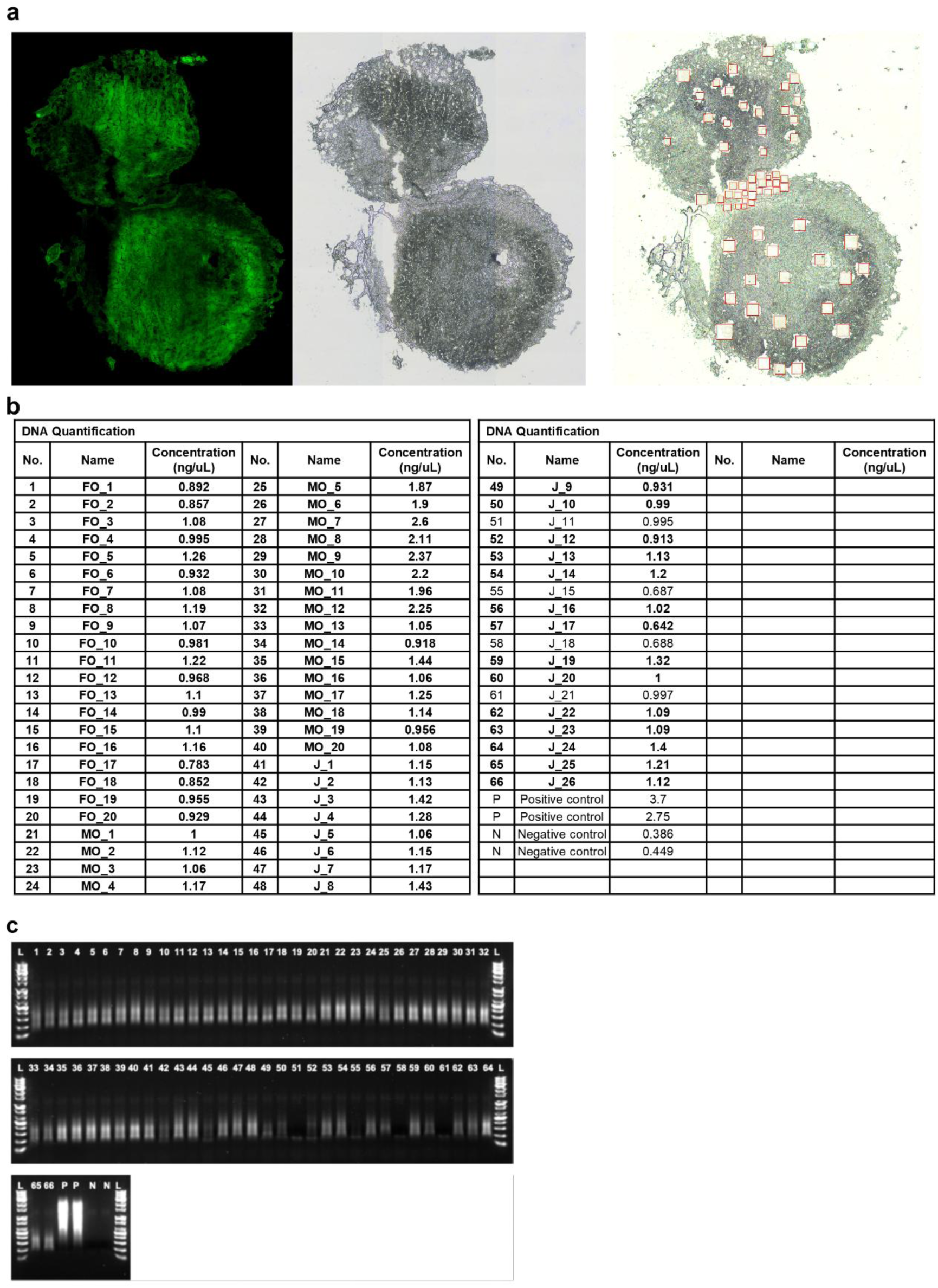
Quality control and sample information for spatially resolved transcriptomic analysis. **(a)** Three regions were analyzed: MO, FO, and junction (J). **(b)** A total of 66 ROIs were isolated (20 MO, 20 FO, and 26 J), of which 62 ROIs (20 MO, 20 FO, and 22 J) passed QC. **(c)** QC pass was determined based on the presence of a smeared band pattern in the electrophoresis profile. Sequencing was performed on 30 ROIs in total (10 from each group). RNA-seq quality assessment results, including RNA concentration and electrophoresis profiles, are shown in the same figure.

**Extended Data fig. 9.**
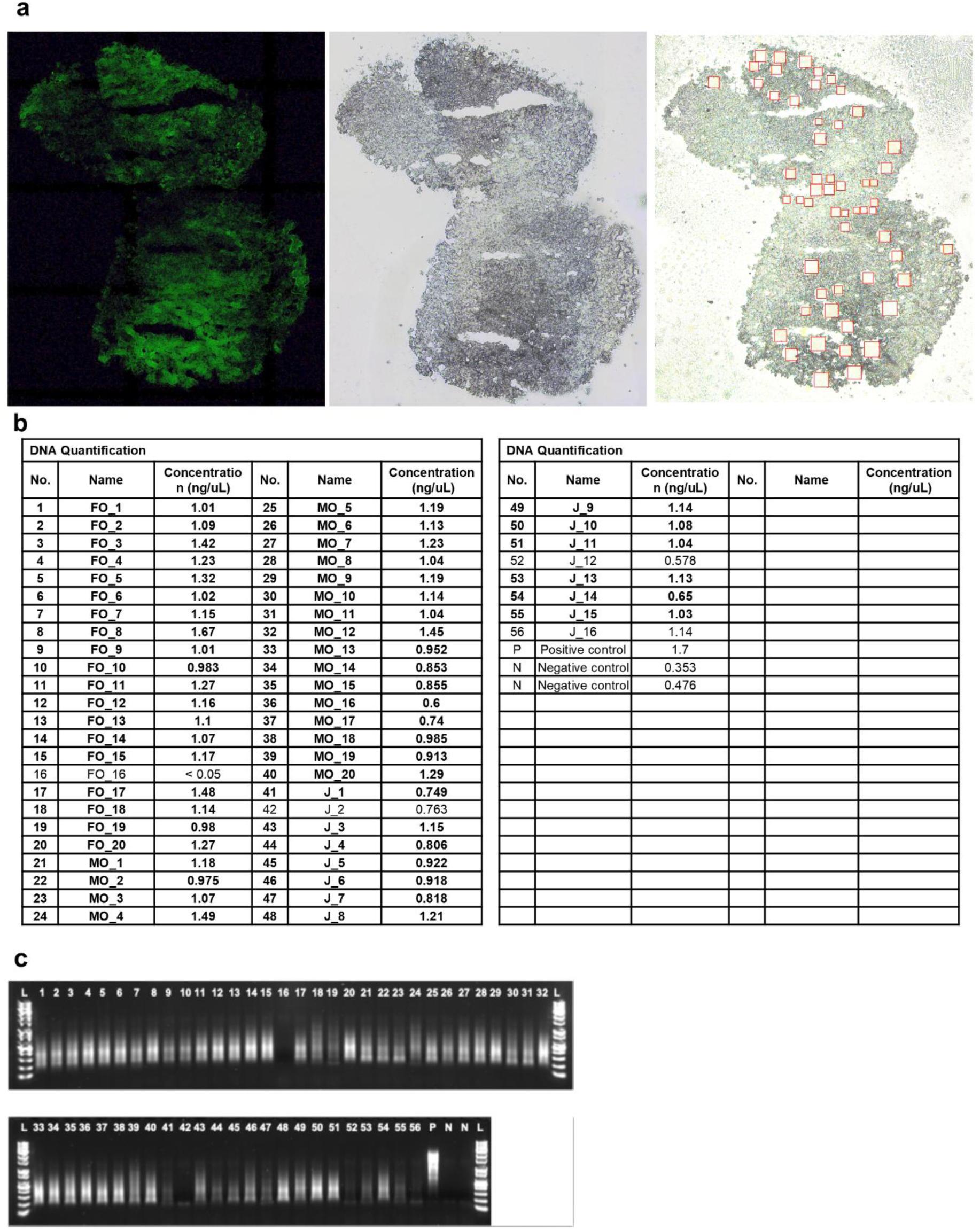
Quality control and sample information for spatially resolved transcriptomic analysis. **(a)** Three regions were analyzed: Three regions were analyzed: MO, FO, and junction (J). **(b)** A total of 56 ROIs were isolated (20 MO, 20 FO, and 16 J), of which 52 ROIs (20 MO, 19 FO, and 13 J) passed QC. **(c)** QC pass was determined based on the presence of a smeared band pattern in the electrophoresis profile. Sequencing was performed on 30 ROIs in total (10 from each group). RNA-seq quality assessment results, including RNA concentration and electrophoresis profiles, are shown in the same figure.

**Extended Data fig. 10.**
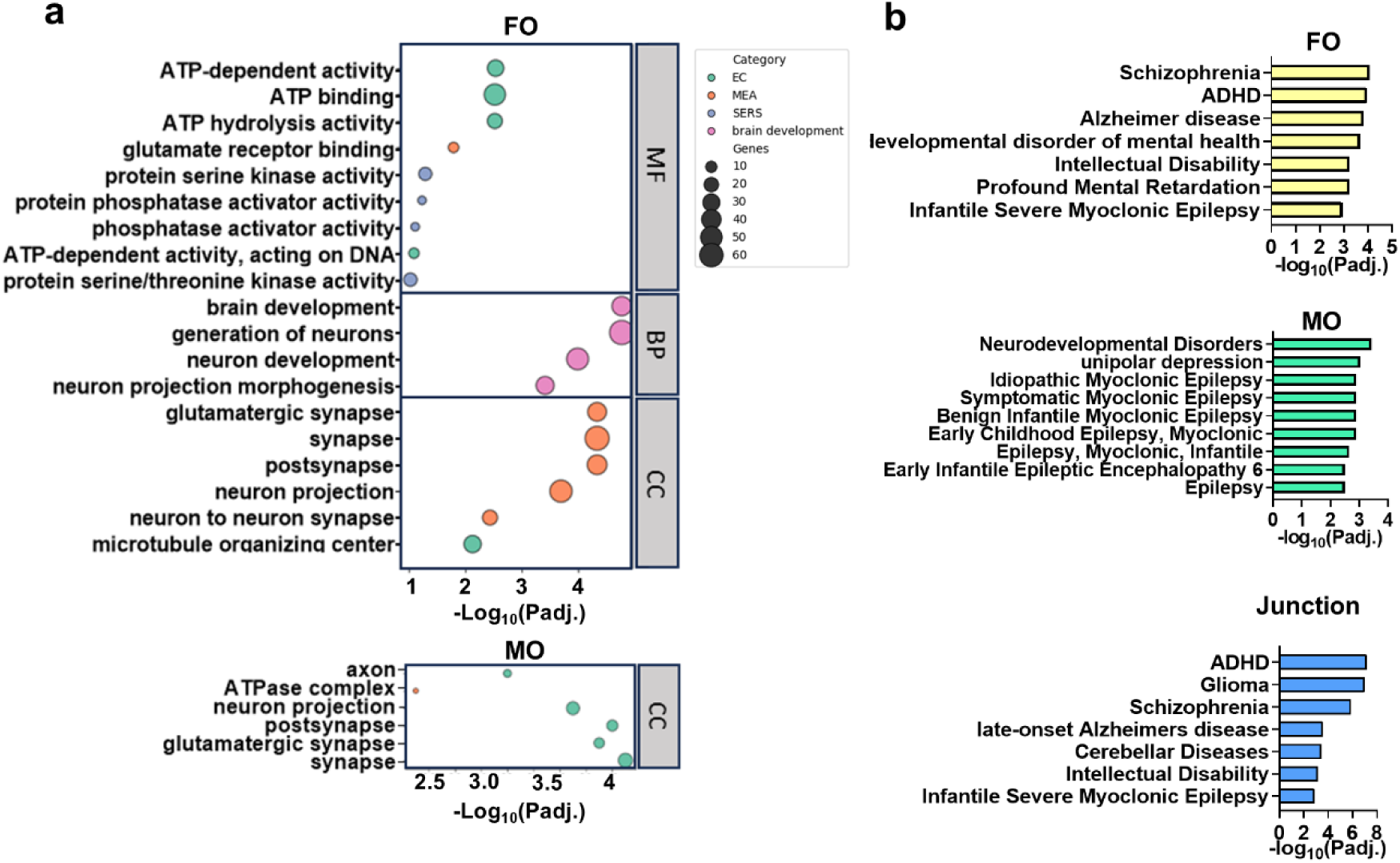
Region-specific gene expression differences revealed by multi-regional spatial transcriptomics in assembloids, highlighting alterations in synaptic function. **(a)** Gene ontology enrichment analysis showing region-specific expression profiles. MF; molecular function, BP; biological process, CC; cellular component. **(b)** Disease ontology enrichment analysis across spatial regions, demonstrating region-specific associations with neurological and neurodevelopmental disorders.

